# Desmin p.R406W mutation is associated with arrhythmias through structural and electrophysiological remodeling

**DOI:** 10.64898/2026.08.05.742729

**Authors:** Michelle Geryk, Thomas Stervinou, Martin Bouaud, Bastien Cimarosti, Jérôme Montnach, Agnès Tessier, Charlène Jouve, Pierre Lindenbaum, Florence Kyndt, Alice Boissard, Cécile Henry, Mélèze Hocini, Sabrina Batonnet-Pichon, Benjamin Lauzier, Guillaume Lamirault, Francois Guillonneau, Jean-Sébastien Hulot, Isabelle Baró, Nathalie Gaborit, Hervé le Marec, Michel Haissaguerre, Vincent Probst, Jean-Jacques Schott, Jean-Baptiste Gourraud, Flavien Charpentier

## Abstract

**Background and Aims:** Mutations in the desmin (*DES*) gene cause a variety of cardiomyopathies associated with arrhythmias, yet the electrophysiological consequences of these variants remain largely uncharacterized. The aim of this study was to investigate the pathogenic mechanisms of the *de novo DES* p.R406W variant, which was identified in a 9-year-old patient who suffered from severe ventricular arrhythmias and sudden cardiac death without overt structural heart disease.

**Methods:** Human induced pluripotent stem cell-derived cardiomyocytes (hiPSC-CMs) carrying the *DES* p.R406W variant (including the patient’s line) were compared to isogenic controls. Action potentials (AP) of hiPSC-CMs were recorded using patch-clamp. Furthermore, 3D engineered heart tissues (EHTs) were generated from hiPSC-CMs and their APs were recorded with sharp microelectrodes. Analytical techniques also included transmission electron microscopy (TEM) and integrated transcriptomic and proteomic profiling. Finally, a heterozygous knock-in (KI) mouse model carrying the *Des* p.R405W ortholog was evaluated through surface ECG, echocardiography and *ex vivo* cardiac optical mapping.

**Results:** The *DES* p.R406W mutation prolonged AP duration in IM-R406W hiPSC-CMs and EHTs vs Control ones. Multi-omics analysis of EHTs revealed a dysregulation of genes and proteins involved in contractile function, cell adhesion, and electrical activity. TEM imaging revealed changes in Z-disc architecture in mutant tissues. Twenty-week-old *Des* p.R405W KI mice exhibited ventricular conduction slowing (prolonged QRS) and a high susceptibility to ventricular tachyarrhythmias, likely due to reentrant mechanisms. Mild hypertrophy was also observed, but only in females.

**Conclusion:** The *DES* p.R406W variant is highly pathogenic, causing electrical and structural remodeling of the myocardium. This study highlights the effectiveness of hiPSC-CMs and EHTs in recapitulating the clinical phenotype of desminopathy, providing a platform for investigating the mechanisms of early-onset cardiac arrhythmias and SCD.

## Introduction

Desmin (OMIM # 125660) is a type Ⅲ muscle-specific intermediate filament (IF) found in cardiac, smooth, and skeletal muscle cells. It is also the most abundant IF in cardiomyocytes (Capetanaki et al. 2007; Tsikitis et al. 2018). Desmin filaments traverse the cell, forming a web that interconnects various parts of the cardiomyocyte, including organelles such as the mitochondria and the nucleus. This ensures their correct location and position within the cell (Capetanaki et al. 2007). Desmin also attaches to Z-discs, interconnecting them laterally and anchoring the contractile apparatus to the intercalated discs (IDs) and costameres (Tsikitis et al. 2018; Hol and Capetanaki 2017; Agnetti et al. 2022). These complex connections contribute to the structural and functional integrity of cardiomyocytes, enabling them to perform their highly demanding function.

Desmin has non-alpha helical N-terminal head and C-terminal tail that flank four alpha helical segments (1A, 1B, 2A, and 2B), which are separated by three linkers (L1, L12, and L2) (Hol and Capetanaki 2017; Agnetti et al. 2022). Mutations in the desmin gene (*DES*) have been identified in all domains. Patients with *DES* mutations have been reported to have skeletal and/or cardiac myopathy, so called desminopathy (OMIM# 601419). Cardiomyopathy in these patients ranges from hypertrophic cardiomyopathy (HCM), restrictive cardiomyopathy (RCM), dilated cardiomyopathy (DCM), with reports of left ventricle non-compaction (LVNC), to arrhythmogenic cardiomyopathy (ACM), with varying symptoms from one patient to the next (Bermúdez-Jiménez et al. 2018; Tsikitis et al. 2018; Kulikova et al. 2021). Age of onset, disease progression, symptoms and cardiomyopathy type differ in unrelated patients carrying the same mutation. This adds to the complexity of desminopathy. The consequences of desmin mutations have been studied using various *in vivo* and *in vitro* models to uncover the underlying mechanisms of disease onset and progression (Herrmann et al. 2020; Goudeau et al. 2006; Bär et al. 2005). Mitochondria (Hovhannisyan et al. 2024; Smolina et al. 2020, 2014; Kubánek et al. 2020), IDs (Herrmann et al. 2020), and Z-disc (Milner et al. 1996; Mavroidis et al. 2008) organization and function have been reported to be significantly affected by desmin mutations. However, although desminopathies have been associated with cardiac conduction blocks and arrhythmias in patients (Geryk and Charpentier 2024), the effect of these mutations on the electrophysiology of cardiomyocytes remains uncharacterized.

In the present study, we investigated the electrophysiological, transcriptomic and proteomic consequences of the *DES* variant p.R406W (c.1216C>T; *DES*^R406W^) located in segment 2B, a highly conserved region across species (Olivé et al. 2004), which was identified in a young female patient presenting with persistent, non-ischemic ST-segment depression on her electrocardiogram (ECG) and severe ventricular arrhythmias, resulting in sudden cardiac death (SCD). To this end, we first used cardiomyocytes derived from human induced pluripotent stem cells (hiPSC-CMs) carrying the *DES*^R406W^ variant (patient’s and isogenic mutant knock-in cell lines) or not (control, unrelated individual’s and patient’s corrected, isogenic control cell lines; *DES*^R406^), which were cultured either as two-dimensional monolayers or three-dimensional engineered heart tissues (EHT). We next investigated the effects of the mutation on a desmin knock-in (KI) mouse carrying the desmin p.R405W mutation (*Des*^R405W^), the ortholog of the human *DES*^R406W^ variant, by performing ECG recordings and *ex vivo* cardiac optical mapping experiments.

## Methods

### Ethical issues

The study was conducted according to the principles set forth under the Declaration of Helsinki (1989) and European guidelines for clinical and genetic research. Institutional review board approvals of the study were obtained before the initiation of patient enrolment. Informed written consent was obtained from each individual who agreed to participate in the clinical and genetic study.

### Genetic investigation

Whole-genome sequencing was performed on the case index and her 2 parents. Genomic DNA (1.1 µg) was used to prepare PCR-free libraries using the Illumina TruSeq DNA Library Kit and sequenced on a HiSeq X5 platform (Illumina) to generate 150 bp paired-end reads. Reads were aligned to the human reference genome (hg19 or hg38) using BWA-MEM, and variants were called with GATK HaplotypeCaller following the Broad Institute’s Best Practices (https://gatk.broadinstitute.org/hc/en-us/sections/360007226651-Best-Practices-Workflows). The mean sequencing depth exceeded 30×, with >90% genome coverage at ≥10×.

### Studies on human induced pluripotent stem cell-derived cardiomyocytes (hiPSC-CMs)

#### Obtention and maintenance of hiPSCs

The four hiPSC lines (PT-R406W, IC-R406, Control, and IM-R406W) used in the current study have been previously described (Geryk et al. 2024; Girardeau et al. 2022). Briefly, the mutant patient-derived line (PT-R406W) and the non-mutant control clone (Control) were generated from peripheral blood mononuclear cells (PBMC) obtained from the patient and an unrelated healthy donor, respectively. The two isogenic lines, namely the patient corrected line (isogenic control; IC-R406) and the control-mutated line (isogenic mutant; IM-R406W), were subsequently generated using CRISPR/Cas9 technology to correct the mutation in the PT-R406W line and introduce the mutation in the Control line, respectively. The success of gene editing was verified using Sanger sequencing and whole genome sequencing (WGS).

All hiPSC lines were maintained at 37°C, 5% CO_2_, 21% O_2_ in StemMACS^TM^ iPS Brew XF Medium (Miltenyi Biotec, Bergisch Gladbach, Germany) on culture plates coated with Matrigel® hESC-Qualified Matrix (0.05 mg/mL, Corning, NY, USA). At 80% confluency, the hiPSCs were passaged using Gentle Cell Dissociation Reagent (STEMCELL^TM^ Technologies, Vancouver, Canada).

#### Cardiac differentiation of hiPSCs

Step specific directed cardiac differentiation of hiPSCs was performed using a modified version of a previously described protocol (Calloe et al. 2022; Treat et al. 2019). Briefly, differentiation was initiated once hiPSCs reached 90% confluency (day 0) by culturing the cells in RPMI1640 medium (Thermo Fisher Scientific, Waltham, MA, USA) supplemented with B27 (without insulin, Thermo Fisher Scientific), 6 µM CHIR99021 (Tocris Bioscience, Bristol, UK), 10 ng/mL recombinant human/mouse/rat activin A protein (R&D Systems, Minneapolis, MN, USA) and 50 µg/ml L-ascorbic acid (Sigma-Aldrich). The medium was renewed at day 1. On days 2, 4, and 6, the culture medium was changed using RPMI1640 supplemented with B27 (without insulin), 5 µM XAV 939 (Hello Bio, Bristol, UK), and 10 µM KY 02111 (Hello Bio). The medium was replaced with RPMI1640 + B27 (with insulin) and L-ascorbic acid on days 8 and 10 followed by a glucose depletion medium composed of RPMI1640 without glucose (Thermo Fisher Scientific) supplemented with B27 (with insulin) and 4 mM sodium L-lactate (Sigma-Aldrich). Cells were kept in depletion medium from days 10-14 and were subsequently changed to maturation medium containing RPMI1640, B27 (with insulin), 1 µM dexamethasone (Cayman Chemical Co, Ann Arbor, MI) and 20 ng/ml 3,3′,5′-Triiodo-L-thyronine sodium salt (T_3_, Sigma-Aldrich). The maturation medium was changed every other day until day 30, after which RPMI1640 + B27 (with insulin) was used. Cardiomyocytes started to spontaneously contract between days 8 and 10.

#### Generation of engineered heart tissues (EHTs) from hiPSC-CMs

EHTs were made from hiPSC-CMs (at least 1×10^6^ cells/EHT) between days 14-17 of differentiation as described previously (Lam et al. 2019; Mannhardt et al. 2017; Ronaldson-Bouchard et al. 2019). Briefly, hiPSC-CMs were dissociated using TrypLE 1x (7 minutes, 37°C, Thermo Fisher Scientific) and centrifuged twice for 3 minutes at 200 g before being resuspended in an EHT mastermix, fibrinogen (200 mg/mL, Sigma-Aldrich) and 10 μl Matrigel® Basement Membrane Matrix (Corning). The EHT mastermix was a combination of sterile filtered 2x DMEM solution [2 ml of heat-inactivated horse serum (Thermo Fisher Scientific), 6 ml of sterile water, and 2 mL of 10x DMEM (134 mg/ml, Thermo Fisher Scientific) and NKM solution (8.9 ml DMEM (Thermo Fisher Scientific) combined with 1 m1 FBS and 100 μl of L-glutamine (Thermo Fisher Scientific)]. For every EHT, 100 μl of the cell-containing solution was briefly mixed with 3 μl of thrombin (100 U/mL, Sigma-Aldrich) and pipetted into a pre-set 2% agarose mold created using a Teflon spacer (DiNAQOR, Hamburg, Germany) in a 24-well plate containing a silicone rack (DiNAQOR). After an initial 2-hour incubation period at 37°C, the silicone racks were gently removed from the molds and added to a new 24-well plate containing a culture medium composed of 50% RPMI1640 + B27 (with insulin), 50% EBM-2 + EGM-2 (Lonza, Basel, Switzerland), 1% penicillin/streptomycin (Thermo Fisher Scientific) with 13.2 mg/l of aprotinin (Sigma-Aldrich), 20 ng/ml of T_3_ hormone, and 1 µM of dexamethasone and placed back into the incubator. This medium was changed every day until day 30. Subsequently, EHTs were switched to a medium containing 50% RPMI1640 + B27 (with insulin), 50% DMEM F-12 + B27 (with insulin), 1% penicillin/streptomycin, and 6.6 mg/L of aprotinin. Medium was changed every other day until needed.

#### Action potential recordings of hiPSC-CMs

The action potentials (AP) of isolated hiPSC-CMs were acquired at 35-37°C using the amphotericin-B perforated-patch configuration of patch clamp as previously described (Al Sayed, Canac, et al. 2021; Al Sayed, Jouni, et al. 2021) with an Alembic VE-2 (Alembic Instruments, Montreal, QC, Canada) or Axopatch 200A amplifier controlled by Axon pClamp 10.6 software through an A/D converter (Digidata 1440A Molecular Devices, San Jose, CA, USA). hiPSC-CMs were superfused with a modified Tyrode solution containing (in mM): NaCl, 130; KCl, 4; HEPES, 10; glucose, 5; MgSO_4_, 1.2; NaH_2_PO_4_, 1.2; CaCl_2_, 1; pH 7.4 (with NaOH). Borosilicate glass pipettes (1.5-3 MΩ of tip resistance, Sutter Instrument, Novato, CA, USA) were pulled on a horizontal puller (P97, Sutter Instruments) and filled with an intracellular solution containing (in mM): NaCl, 5; KCl, 20; HEPES, 5; K-Gluconate, 125; pH 7.2 (with KOH). Amphotericin-B (Sigma) was added extemporarily to the intracellular solution (0.2-0.4 μg/mL). APs of spontaneously beating hiPSC-CMs were recorded first. Since hiPSC-CMs have a limited I_K1_, artificial I_K1_ was injected using dynamic patch-clamp (Meijer van Putten et al. 2015; Wilders 2006). A custom-made software that runs on RT-Linux allowed injection of I_K1_ through an A/D converter (PCI-6221, National Instruments, Austin, TX, USA) connected to the voltage output and the current command of the patch-clamp amplifier. By controlling I_K1_ amplitude, the membrane potential was set to -90 mV. Cells were paced using a 1-ms square depolarizing current pulsed at a pacing cycle length (PCL) of 1000 ms and 700 ms. APs were analyzed using a custom R script that analyzed individual APs from files created in Clampfit 10.6 during acquisition of either spontaneous or paced APs. The AP maximum diastolic potential (MDP), overshoot, amplitude, maximum upstroke velocity (dV/dt_max_), and duration (APD) at different levels (30%, 50%, 90%) of full repolarization were measured in spontaneously beating and I_K1_-clamped hiPSC-CMs. Data from four to five consecutive APs were averaged. In addition, peak-to-peak interval (beating rate) was measured in spontaneously beating hiPSC-CMs.

#### Action potential recordings of EHTs

APs were recorded from spontaneously beating EHTs using high-resistance sharp glass microelectrodes (40-60 MΩ tip resistance; Harvard Apparatus) pulled using a horizontal puller (P97, Sutter Instrument) and filled with 3 M KCl. The EHTs were superfused in a custom-made, 3D-printed chamber, with the same Tyrode solution used for the patch-clamp experiments, and bubbled with 100% O_2_. Microelectrodes were connected to an amplifier (VF-102, BioLogic, Seyssinet-Pariset, France) and voltage traces were visualized on an oscilloscope (Tektronix, Beaverton, OR, USA) and digitized with an analogic/digital converter (PowerLab C, ADInstruments, Dunedin, New Zealand) at a sampling rate of 20 kHz for recording with LabChart 8 Pro software (AD Instruments). All AP recordings were performed at 36-37°C. Intervals between APs, MDP, AP amplitude, overshoot, dV/dt_max_ and APDs were analyzed as for isolated hiPSC-CMs. Data from five consecutive APs were averaged.

#### Transcriptomic and proteomic studies

##### Sample collection

For all four hiPSC lines, samples were collected from 5 to 8 independent EHTs on day 45 of differentiation. EHTs were washed four times with DPBS solution without Ca^2+^and Mg^2+^ at 4°C, removed from the silicone rack and then cut into two pieces. For every EHT, one half was flash frozen in a Cryotube (Thermo Fisher Scientific) in liquid nitrogen and stored at -80°C for proteomic applications and the second half was resuspended in RA1 (Macherey-Nagel, Dueren, Germany) and stored at -80°C for transcriptomic applications.

##### Data generation

For 3’RNA-Sequencing, total RNA was extracted using the NucleoSpin RNA kit (Macherey-Nagel) according to manufacturer’s instructions and it’s quality assessed by NanoDropTM 1000 Spectrophotometer (Thermo Fisher Scientific). 3’RNA libraries were prepared by GenoBiRD core facility as previously published (Charpentier et al. 2021) and sequenced on a NovaSeq 6000 Sequencing System (Illumina, San Diego, CA, USA).

For proteomics (non-targeted LC-MS/MS performed at the Prot’ICO facility), EHTs were quickly thawed and proteins were concomitantly extracted and denatured using 200 µL of 0.1% Rapigest® SF acid-labile detergent (Waters), 5 mM DTT and 50 mM ammonium bicarbonate, at 95°C for 30 min. Thiol residues were thus chemically reduced and subsequently protected by thiomethylation in 10 mM MMTS (Sigma) for 10 min at 37°C. Samples were cooled to room temperature before adding 1 µg trypsin (Porcine, sequencing grade from ABSciex); for 50 µg of protein, incubated at 37°C overnight. Peptides were then cleared by centrifugation, desalted using Oasis HLB solid phase extraction device (Waters). Eluates were dried in a vacuum centrifuge concentrator (Thermo Fisher Scientific), resuspended in 25 µL of 10% Acetonitrile (ACN) and 0.1% Formic Acid (FA). The equivalent of 200 ng of peptides was injected after microBCA peptide assay (Thermo Fisher Scientific).

Each sample was injected and separated on a C_18_ reverse phase column (Aurora series 1.6 µm particles size, 75-µm inner diameter and 25-cm length from IonOptics) using a NanoElute LC system (Bruker, Billerica, MA, USA). Eluate flow was electrosprayed into a timsTOF Pro 2 mass spectrometer (Bruker) for the 60-min duration of the hydrophobicity gradient ranging from 99% of solvent A containing 0.1% FA in milliQ-grade H_2_O to 40% of solvent B containing 80% ACN plus 0.1% FA in mQ-H_2_O. The mass spectrometer acquired data throughout the elution process and operated in data-independent analysis with PASEF-enabled method using the TIMS-Control software (Bruker). Samples were injected in batch replicate order to circumvent possible technical biases.

##### Data availability

The raw data generated by DIA-PASEF and extracted proteomics data have been deposited to the ProteomeXchange Consortium via the PRIDE (Perez-Riverol et al. 2025) partner repository with the dataset identifier PXD081882.

##### Data analysis

For transcriptomics, demultiplexing, alignment on GRCh38 reference genome, counting steps, normalization and log-transformation of expression matrices were conducted with the Snakemake pipeline developed by the GenoBiRD core facility (https://bio.tools/3SRP) (Charpentier et al. 2021). Genes with significant expression variation between *DES*^R406^-EHTs (Control and IC-R406 lines) and *DES*^R406W^-EHTs (PT and IM-R406W lines) were identified with DESeq2 (RRID:SCR_015687) (Love et al. 2014). Benjamini-Hochberg-corrected p-value < 0.01 and |Log2(FoldChange) |>|1| was set as detection threshold. Principal Component

Analysis (PCA) was performed with the R package FactoMineR (RRID:SCR_014602) (Lê et al. 2008) on mean-centered and log-transformed data matrices. Heatmaps of differentially expressed genes (DEGs) were displayed using the R package ComplexHeatmap (RRID:SCR_017270) (Gu 2022) on scaled and centered log-transformed matrices. Volcano plots of DEGs were generated using the R package EnhancedVolcano (RRID:SCR_018931) (Blighe et al. 2018). Functional annotation [Gene Ontology (GO) and Disease Ontology (DO) enrichment] was performed using the R package ClusterProfiler (RRID:SCR_016884) (Xu et al. 2024).

For proteomics, the raw data were normalized and analyzed using Spectronaut 18.0.23 (Biognosys) in DirectDIA+ mode, which modelized elution behavior, mobility and MS/MS events based on the Uniprot/Swissprot sequence 2022 database of human proteins. Protein identification false discovery rate (FDR) was restricted to 1% maximum, with a match between runs option enabled, and inter-injection data normalization. The enzyme’s specificity was trypsin’s with up to two missed cleavage sites. The precursor and fragment mass tolerances were set to 15 ppm. Oxidation of methionines was set as variable modifications while thiol groups from cysteines were considered completely modified by thiomethylation. A minimum of two ratios of peptides was required for relative quantification between groups. Protein quantification analysis was performed using Label-Free Quantification (LFQ) intensities. Proteins were only considered for further analysis when identified in ≥ 75% of samples. Missing values were imputed with nearest neighbor averaging (KNN) using the impute.knn function from the IMPUTE (RRID:SCR_009245) R package (Howie et al. 2009). The proteins LFQ values were normalized using quantile normalization then log2-transformed and stored in a matrix.

PCA was performed with the R package FactoMineR (RRID:SCR_014602) (Lê et al. 2008) on the entire normalized and log-transformed matrix. Proteins with significant expression variation between *DES*^R406^-EHTs and *DES*^R406W^-EHTs were identified with a Limma test from the R package LIMMA (RRID:SCR_010943) (Ritchie et al. 2015) and a Benjamini-Hochberg-corrected p-value < 0.05 based on a log2-transformed matrix with missing values imputed.

#### Transmission electron microscopy

EHTs were fixed on day 47 (+/-2 days) in 2.5% glutaraldehyde in phosphate buffer (DPBS without Ca^2+^ and Mg^+^) at pH 7.4 for 24 hours at 4°C. The EHTs were then removed from the silicone pillars and transferred into a phosphate buffer containing 81% Na_2_HPO_4_ 67.04 mM and 19% NaH_2_PO_4_ 65.2 mM at a pH 7.4 and stored at 4°C prior to being sent to the SCIAM facility in Angers (France). There, samples were post-fixed in 1% osmium tetroxide/1% potassium ferrocyanide in water for 1h30 at room temperature. Then, the samples were washed three times by deionized water and subsequently dehydrated in a graded series of 35% (2×15 min), 50% (2×15 min), 70% (2×15 min), 95% (2×15 min) and 100% ethanol (2×30 min), followed by propylene oxide (2×15 min). After being infiltrated with a 1:1 propylene oxide and epoxy resin (Epon™ 812) overnight, the samples were embedded in 100% epoxy resin, which was then left to polymerize for 24 h at 37°C, followed by 24 h at 45°C and 48 h at 60°C. Ultra-thin sections (60 nm) were cut from each sample using Leica UC7 ultramicrotome (Leica microsystems, Wetzlar, Germany) and deposited onto copper grids. Sections were the stained with 3% uranyl acetate in 50% ethanol for 15 min, washed three times with deionized water, stained with 3% lead citrate and washed again three times with deionized water. The samples were then left to dry and subsequently examined using the JEOL JEM-1400 electron microscope (JEOL, Tokyo, Japan) under 120 keV.

#### Micropatterning and immunostaining

Frozen hiPSC-CMs were thawed on D29 and plated onto Matrigel coated 12-well plates with 1/1000 ROCK inhibitor. The culture medium was changed every other day using RPMI1640 containing B27 (plus insulin). Seven days later (D36), cells were dissociated and 250 000 hiPSC-CMs were seeded onto custom designed micropatterned 18 mm coverslips (4D Cell, Montreuil, France). The micropatterned lines were 30 μm wide and 100 μm apart. Similarly, the following days, the medium was changed every other day using RPMI1640 with B27 (plus insulin). One-week later (D42), the seeded hiPSC-CMs were fixed with 4% paraformaldehyde for 10 min at RT, permeabilized with 0.5% Triton X-100 in PBS for 15 min and blocked with 2% BSA in PBS for 1 hour. Cells were incubated overnight at 4°C with primary antibodies diluted in 1% BSA in PBS, washed three times with PBS and incubated with suitable secondary antibodies and DAPI for 1 hour at room temperature. Desmin primary antibody (REF: HPA018803, Sigma) and the desmoplakin primary antibody (REF: 61024, Thermo Fisher Scientific) were both diluted at 1:100. Secondary antibodies were diluted at 1:1000 (goat anti-rabbit 546 REF: A11010, Thermo Fisher Scientific; goat anti-mouse 488 REF: A10680, Thermo Fisher Scientific). Immunostainings were examined using an inverted Eclipse Ti2 fluorescence microscope (Nikon) using Nikon Standard software.

### Mouse Studies

#### Study approval – animal ethics

All experiments on mice were performed at the animal facility of Nantes University Health Research Institute (UTE - IRS-UN). They were approved by the regional ethics committee on animal experimentation and authorized by the French Ministry of National Education, Higher Education and Research, according to the Directive 2010/63/EU of the European Union (agreement APAFIS #43783-2023042420021419v6).

#### Desmin p.R405W knock-in mouse model

In this study, only KI mice that were heterozygous for the Des-p.R405W mutation (Hakibilen et al. 2022; Herrmann et al. 2020) and their wild-type littermates were investigated at 10 and 20 weeks of age. Genotyping of the animals was performed on tail fragments with Thermo Scientific^TM^ Phire^TM^ Tissue Direct PCR kit following the manufacturer recommendations. Genotypes were determined with primers 5′-CTGGAGGAGGAGATCCGACA-3′ and 5′-GGCCCTCGTTAATTTTCTGC-3′ (Hakibilen et al. 2022). All experiments have been performed on both male and female mice.

#### ECG recording and analysis

Mice were anesthetized with isoflurane: 3% in air in an induction chamber for 2-3 minutes followed by 1.5-2% in air by mask throughout the recording. Body temperature was maintained at 37°C using a retro-controlled heating pad (Harvard Apparatus, USA). A six-lead surface ECG was recorded with 25-gauge subcutaneous electrodes connected to a computer through an analog-digital converter (IOX 1.585, EMKA Technologies, Fr). This allowed for real-time monitoring and subsequent offline analysis of the ECG data using ECG Auto v3.2.0.2 (EMKA Technologies). ECG parameters were measured on lead I. The QRS interval was calculated from the onset of the Q wave to the point where it intersects the isoelectric line and the ST segment. We also determined the amplitude of the Q, R, S and J (positive wave following the QRS complex) waves (Boukens et al. 2014). The criteria for measuring RR, PR and QT intervals, along with P-wave duration, have been previously documented (Royer et al. 2005). Validation of the measurements was performed by an operator blinded to the genotype on extracted ECG complexes.

#### Optical mapping of [Ca^2+^]_i_

Optical mapping experiments were performed by operators blinded to the genotype. Twenty-one-week-old mice were heparinized (600 IU/kg i.p.) and euthanized by cervical dislocation. The heart was quickly excised and the aorta was cannulated in ice-cold Ca^2+^-free modified Krebs-Henseleit (KH) solution containing (in mmol/L): NaCl, 116; NaHCO_3_, 27; NaH_2_PO_4_, 0.35; KCl, 5; MgSO_4_ (7 H_2_O), 1.1; glucose, 10. The heart was then mounted onto a Langendorff perfusion system and placed into the bath of a dual signal optical mapping system (MappingLab Ltd, Oxford, UK). The heart was then perfused with a modified KH solution (in mmol/L: NaCl, 116; NaHCO_3_, 27; NaH_2_PO_4_, 0.35; KCl, 5; MgSO_4_ (7 H_2_O), 1.1; CaCl_2_, 1.8; Na-pyruvate, 0.2; Na-lactate, 1; glucose, 10; 95.% O_2_/5% CO_2_) at 2 mL/min and 37°C. Two electrodes were placed near the apex of the heart (positive electrode) and the atria (negative electrode) to record a 1-lead extracorporeal ECG (reference electrode pinned in the bath). After 5 min of stabilization, electromechanical uncoupling was obtained with a 10-min perfusion with blebbistatin (5.5 µmol/L; Tocris Bioscience). Then the heart was progressively perfused first with a mix of Rhod-2 AM (StemCell^TM^ Technologies) and Pluronic F-127 (Invitrogen^TM^, Thermo Fisher Scientific) during 10 minutes, then with RH 237 (Invitrogen^TM^, Thermo Fisher Scientific) for 10 additional minutes for simultaneous optical recording of [Ca^2+^]_i_ transients (CaT) and V_m_. The heart was excited with a light emitted at a wavelength of 530 ± 25 nm and bandpass filtered at 511–551 nm (LEDC-2001, MappingLab Ltd). The high-pass filters for Rhod-2 and RH 237 were 585 nm and 700 nm, respectively. Fluorescence signals were recorded with 2 CMOS cameras (OMS-PCIE-2002 Beta, MappingLab Ltd).

After recording spontaneous activity, the heart was paced with a pair of bipolar electrodes positioned at 1 mm from the apex of the heart using a programmable stimulator (VCS-3001, MappingLab). Pacing stimuli were 2-ms square wave currents with an amplitude set at 1.5-2 threshold current. AP and CaT activation patterns, conduction velocity and duration as a function of heart rate were determined with 10-second S1S1 pulse trains at 7, 8, 9 and 10 Hz, separated by at least 30-second intervals. Ventricular effective refractory period (ERP) was determined with an S1S2 protocol. A single extra stimulus (S2) was delivered after 15 consecutive S1 at a basic cycle length of 100 ms (10 Hz) and decremented in 4-ms steps from S1S2 = 70 ms to S1S2 = 20 ms. S1S2 pacing trains were separated by 20-seccond intervals. Finally, a third pacing protocol consisted in a 10-second burst with S1S1 progressively decreasing by 1-ms decrements from 150 ms at the start of the burst to 50 ms at its end. S1S2 and burst pacing protocols were performed under baseline condition and after 5 minutes of perfusion with isoprenaline (0.1 µmol/L).

In some mice, S1S2 and burst pacing protocols triggered ventricular tachyarrhythmias. Only ventricular tachyarrhythmias occurring at the end of the pacing trains were considered. Tachyarrhythmias were categorized into 5 distinct types: single ventricular premature beat (VPB), doublet of VPBs, triplet of VPBs, short ventricular tachycardia (VT; < 15 beats) and sustained VT (≥ 15 beats).

#### Transthoracic echocardiography

Non-invasive transthoracic echocardiography was performed on sedated mice to assess cardiac function *in vivo* using a Vevo 2100 system (40-MHz transducer; Visualsonics, Toronto, Canada). Mice were anesthetized by inhalation of isoflurane (as for ECG recording) and placed in supine position on a temperature-controller examination table to maintain rectal temperature at 37°C. ECG was monitored via limb electrodes during the whole procedure. Time motion mode recording in parasternal long-axis view allowed to measure various parameters: end-diastolic and end-systolic left ventricular posterior wall thickness (LVPWTd and LVPWTs), end-diastolic and end-systolic left ventricular internal diameter (LVIDd and LVIDs), ejection fraction (EF) and heart rate (HR).

#### Histology

Mice were euthanized by cervical dislocation, and the heart was isolated and washed with PBS, fixed in 4% paraformaldehyde and embedded in paraffin. Serial sections of 5 μm thickness were stained with picrosirius red as previously described (Derangeon et al. 2017). Stained sections were examined with a classic light microscope (Nikon Eclipse Ti2) and pictures were acquired with NIS-Elements software (v4.10, Nikon, Japan). Fibrosis quantification was done with Fiji (Schindelin et al. 2012) on one coronal section per heart.

Delineation of the regions of interest (ROI) was done by an operator blinded to the genotype and quantification was performed automatically on series of sections by a custom-made Fiji program. For each section, results from 2-3 adjacent ROIs covering as much as possible of the right ventricular free wall, septum and left ventricular free wall were averaged.

#### Statistics

Statistical analyses were conducted using Prism software (GraphPad Software, Inc., USA). Statistical significance was evaluated using a Student t-test or Mann-Whitney test for comparing two groups. For comparisons involving more than 1 factor and repeated measures, a two-way ANOVA for repeated measures was applied, followed by Tukey or Sidak test for multiple comparisons. Fisher exact test was used to compare qualitative variables. Statistical tests are specified in the figure and table legends. Statistically significant differences between values were defined as 2-tailed *P*< 0.05.

## Results

### Case study

A 9-year-old female patient was admitted to the hospital after she was successfully resuscitated from SCD. Her medical history was uneventful and with no family history of SCD. A complete clinical evaluation including echocardiography and cardiac computed tomography did not reveal any structural heart disease, including abnormalities of the coronary arteries. In contrast, the patient’s ECG revealed a deep and persistent ST-segment depression in leads I, II, aVL, and V3 through V6 and in association with a coved ST segment elevation in leads III, aVR and V1 (Figure 1A). An exercise test did not trigger any arrhythmia. This ECG pattern persisted over time. She received an implantable cardioverter-defibrillator and was treated with hydroquinidine. Familial screening was performed in all the first-degree relatives of the patient and did not reveal any ECG abnormalities.

**Figure 1.**
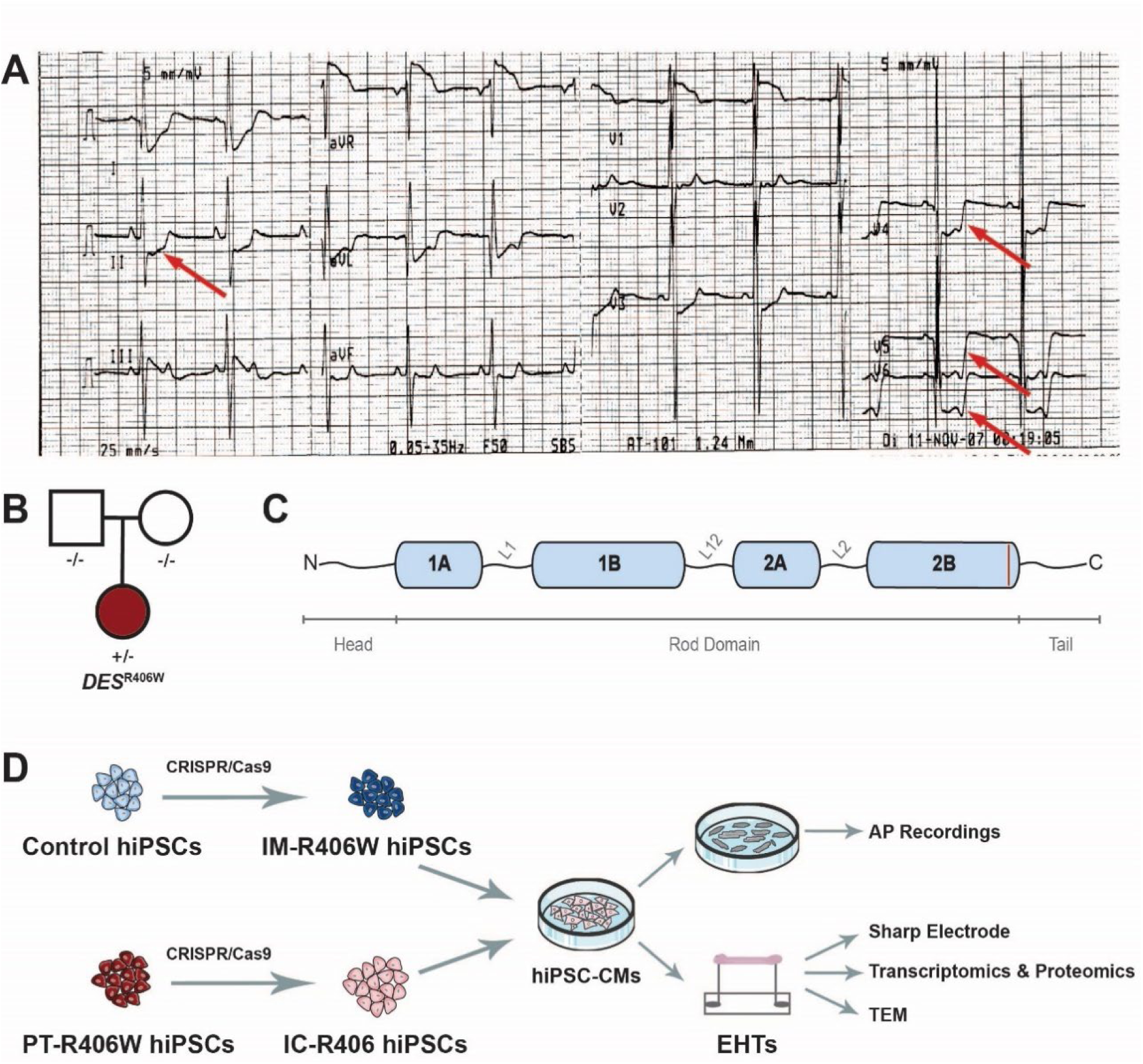
*De novo* heterozygous *DES*^R406W^ variant. **A**. Representative patient ECG demonstrating ST-segment depression (red arrows; leads II, and V4-6). **B**. Family pedigree: circles and squares represent female and male individuals, respectively. Empty symbols represent unaffected family members. The dark red circle represents the affected patient. **C**. Schematic representation of a desmin monomer that is composed of a non-helical N-terminal head and C-terminal tail flanking a rod domain composed of four segments (in blue: 1A, 1B, 2A, and 2B) that are separated by three linkers (L1, L12, and L2). The *DES*^R406W^ variant is located in the terminal part of segment 2B (dark red line). **D.** Schematic overview of the hiPSC lines and techniques used. **Top left.** Control hiPSCs (unrelated to the patient; light blue) were used to generate an isogenic mutant line (IM-R406W; dark blue) containing the *DES*^R406W^ variant using CRISPR/Cas9. **Bottom.** An isogenic control line (IC-R406; light pink) was generated from the patient’s hiPSCs (PT-R406W; dark red) using CRISPR/Cas9. **Middle.** All four cell lines were differentiated into hiPSC derived cardiomyocytes (hiPSC-CMs). **Top right.** hiPSC-CMs from all clones were used for action potential (AP) recordings using the patch clamp technique. **Bottom right.** Engineered heart tissues (EHTs) of all clones were generated from hiPSC-CMs, and were used for transcriptomic and proteomic studies, transmission electron microscopy (TEM), and AP measurements using sharp electrodes. Figure 1D was adapted from Servier Medical Art (https://smart.servier.com), licensed under CC BY 4.0 (https://creativecommons.org/licenses/by/4.0/).

After an initial 18-month period with 2 recurrences of ventricular fibrillation (VF), the patient’s condition deteriorated to at least one VF episode per month. Several antiarrhythmic drugs (including beta-blocker therapy and amiodarone) were tried; first separately and finally combined, without efficacy. Her condition eventually worsened to several episodes of VF per day during several days leading to an alteration of her hemodynamic status. VF episodes were initiated from polymorphic ventricular premature beats (VPBs), mainly originating from the left ventricle (LV). The severity of her condition prompted an electrophysiological study, during which a few Purkinje potentials were detected. This finding may be associated with the persistent conduction defects and ST-segment modification observed on the ECG. Multiple catheter ablations in various VPB locations, namely the left interventricular septum, the right ventricular outflow track, and the LV apex and anterior wall were ineffective, as new electrical storms emerged from different loci, ultimately leading to cardiogenic shock. The patient was placed on extracorporeal membrane oxygenation because of her weakened LV function, and finally transplanted at the age of 12. She has since remained healthy, with a follow-up period of 15 years.

Anatomo-histopathological examination showed that both the right ventricle and focally the LV exhibited increased amounts of epicardial adipose tissue but no overt evidence of fibro-fatty replacement. In the LV, small areas of fibrosis and scarring, which may represent ischemic changes possibly caused by VF, were noted, in addition to ablation lesions. A clear structural substrate for the clinical presentation of the patient was not observed on histology. Evidence for myocarditis, pericarditis, endocarditis or vasculitis was not noted.

### Genetic investigation

Whole genome sequencing was performed on the patient and her siblings, leading to the identification of a *de novo* variant (c.1216C>T) in the *DES* gene (Figure 1B). This variant results in a substitution of the arginine (R) by a tryptophan (W) at position 406 (p.R406W) in segment 2B of the desmin protein (Figure 1C: red line).

### Electrophysiological characterization of hiPSC-CMs

To elucidate the electrophysiological consequences of the heterozygous *DES*^R406W^ mutation, four hiPSC lines were differentiated into hiPSC-CMs: (1) a patient-unrelated control line (Control), which was used to generate (2) an isogenic mutant line (IM-R406W) containing the *DES*^R406W^ variant, (3) the patient’s line (PT-R406W), which was used to generate (4) an isogenic control line (IC-R406) with two *DES*^R406^ wildtype alleles (Figure 1D). The electrophysiological phenotype was evaluated by recording spontaneous (Figure 2A, left panels) and paced action potentials (APs) (Figure 2A, right panels). The beat-to-beat interval of the spontaneously beating hiPSC-CMs did not differ between the cell lines (Figure 2B). There was a significant difference in maximum diastolic potential (MDP) between the two compared groups. Indeed, IM-R406W hiPSC-CMs had a less negative MDP (−52.4 ± 1.3 mV) than Control hiPSC-CMs (−56.8 ± 1.6 mV, p=0.03) (Figure 2C). Similarly, MDP in PT-R406W hiPSC-CMs (−49.3 ± 1.2 mV) was less negative than in IC-R406 hiPSC-CMs (−54.2 ± 1.5 mV, p=0.008). There were no significant differences in AP amplitude between the four lines (Figure 2E).

**Figure 2.**
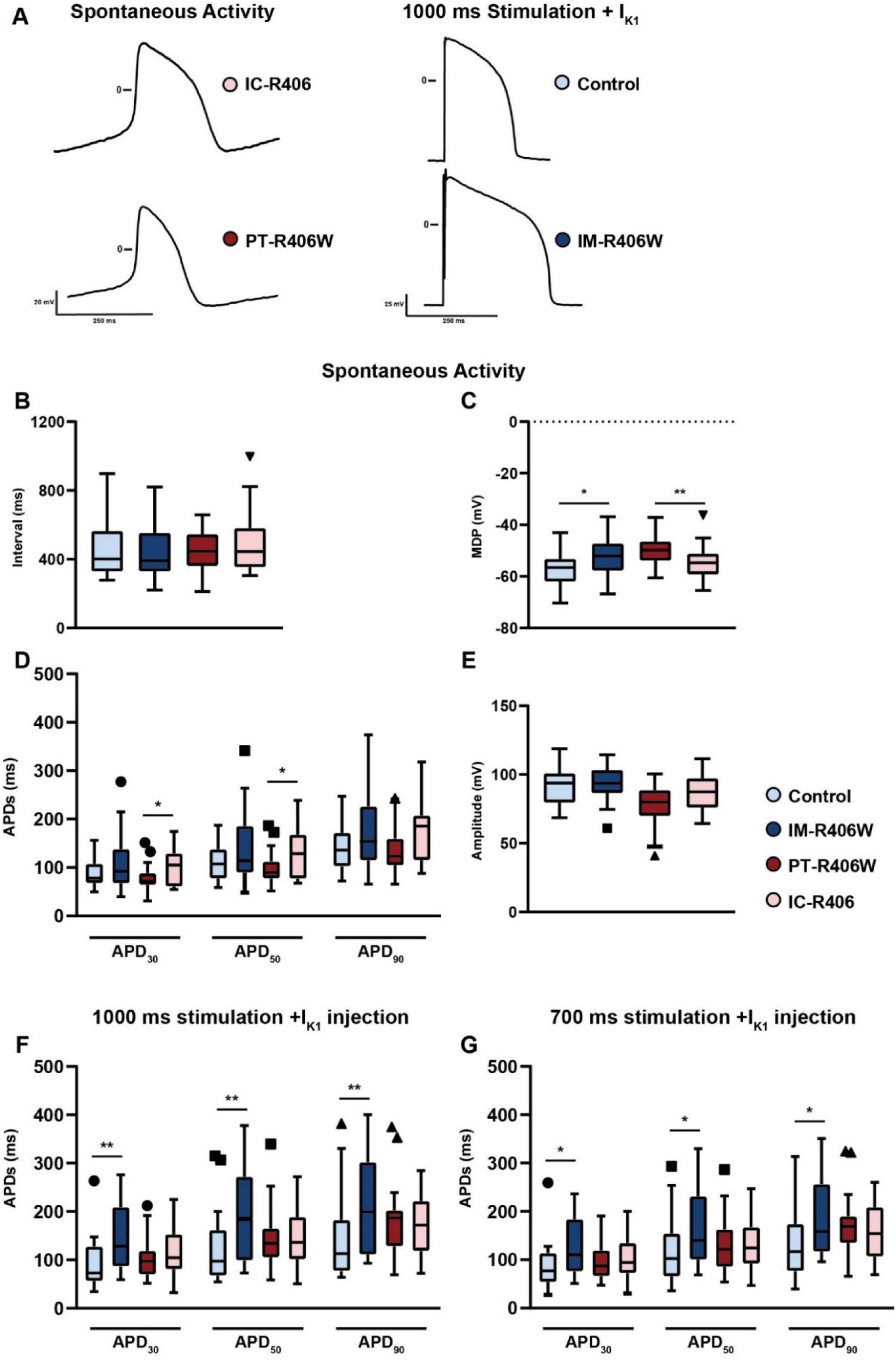
*DES*^R406W^ variant leads to changes in hiPSC-CM APs. Patch-clamp experiments comparing various AP parameters of Control, patient (PT-R406W), and isogenic (mutant; IM-R406W, control; IC-R406) hiPSC-CMs. **A**. Representative APs of spontaneously beating hiPSC-CMs (left) and stimulated (PCL = 1000 ms) hiPSC-CMs with I_K1_ injected using dynamic clamp (right). Beat-to-beat interval (**B**), maximum diastolic potential (MDP; **C**), AP duration (APD) at 30, 50, and 90% of repolarization (APD_30_, APD_50_, APD_90_, respectively; **D**), and amplitude (**E**) of spontaneously beating Control (n=19), IM-R406W (n=25), PT-R406W (n=24), and IC-R406 (n=20) hiPSC-CMs. **F**. APD_30_, APD_50_, APD_90_ of Control (n=22), IM-R406W (n=15), PT-R406W (n=30), and IC-R406 (n=28) hiPSC-CM at PCL of 1000 ms with injected I_K1_. **G**. APD_30_, APD_50_, and APD_90_ of Control (n=21), IM-R406W (n=15), PT-R406W (n=27), and IC-R406 (n=25) hiPSC-CM clones at PCL of 700 ms with injected I_K1_. Mann-Whitney test was used to assess significance between Control versus IM-R406W and PT-R406W versus IC-R406 (*: p < 0.05, **: p < 0.01). Data represented as Tukey box plots.

PT-R406W hiPSC-CMs exhibited shorter APD_30_ and APD_50_ (76.4 ± 5.6 mV, p=0.03, and 98.4 ± 6.9 mV, respectively) than IC-R406 hiPSC-CMs (103.8 ± 8.5 mV and 131.0 ± 11.2 mV, respectively; p=0.04) (Figure 2D). There were no significant differences in the APDs of the Control and IM-R406W lines (Supplementary Table 1).

Due to the lack of intrinsic I_K1_ in hiPSC-CMs, we used the dynamic patch-clamp to inject computed I_K1_ and impose an MDP around -90 mV. The cells were paced at cycle lengths of 1000 and 700 ms. As observed in spontaneously beating cells, the APD_30_, APD_50_ and APD_90_ were longer in IM-R406W hiPSC-CMs (149.5 ± 18.4 mV, 198.5 ± 25.6 mV, and 217.6 ± 26.8 mV, respectively) than in Control hiPSC-CMs (92.2 ± 11.0 mV, 112.3 ± 15.9 mV, and 138.9 ± 18.1 mV, respectively) when paced at 1000 ms (Figure 2F). A similar trend was observed between the two lines at a PCL of 700 ms (Figure 2G): IM-R406W hiPSC-CMs had longer APDs (APD_30_: 128.5 ± 16.2 mV, APD_50_: 168.6 ± 21.3 mV, and APD_90_: 189.8 ± 21.6 mV) than

Control hiPSC-CMs (APD_30_: 85.5 ± 10.9 mV, APD_50_: 112.8 ± 14.5 mV, and APD_90_: 129.2 ± 16.2 mV). In contrast, there were no significant differences in APD between PT-R406W and IC-R406 hiPSC-CMs (Supplementary Table 1).

### Electrophysiological characterization of EHTs

To assess the properties of the *DES*^R406W^ mutation in a three-dimensional model, we generated EHTs from the four hiPSC-CM lines (Figure 1D) and recorded their APs with sharp microelectrodes. Representative APs of the four different EHTs are shown in Figure 3A. No difference in beat-to-beat interval between spontaneously beating EHTs carrying the *DES*^R406W^ mutation (IM-R406W and PT-R406W) and their respective controls (Control and IC-R406, respectively; Figure 3B). There were no significant differences in the overshoot (Figure 3C) or MDP (Figure 3E) between the conditions.

**Figure 3.**
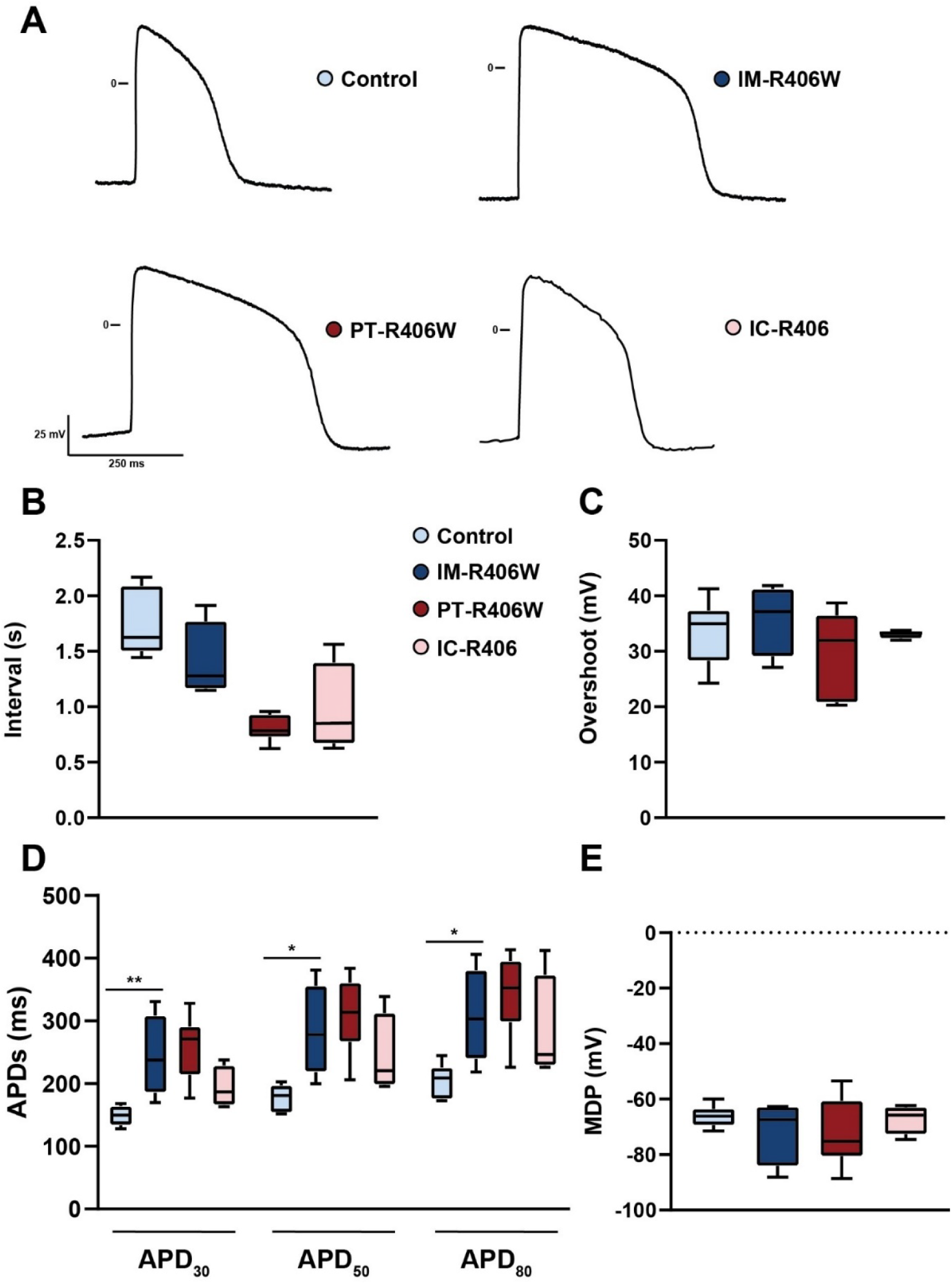
Effects of the *DES*^R406W^ variant on AP parameters of engineered heart tissues (EHTs). **A.** Representative APs of spontaneously beating EHTs. Peak-to-peak interval (**B**), overshoot (**C**), AP duration at 30, 50, and 80% of repolarization (APD_30_, APD_50_, APD_80_, respectively; **D**), and maximum diastolic potential (MDP) **(E**), of spontaneously beating control (n=6), IM-R406W (n=4), PT-R406W (n=7), and IC-R406 (n=4) EHTs. Mann-Whitney test was used to assess significance between Control versus IM-R406W and PT-R406W versus IC-R406 (*p < 0.05, **p < 0.01). Data represented as Tukey box plots.

Moreover, the IM-R406W EHTs had significantly longer APD_30_, APD_50_ and APD_80_ (244.0 ± 33.1 ms, 284.3 ± 37.1 ms, 308.0 ± 38.3 ms, respectively) compared to Control EHTs (149.0 ± 6.6 ms, 177.8 ± 8.7 ms, 205.0 ± 11.1 ms, respectively; Figure 3D). A similar trend, that did not reach significance, was observed between the PT-R406W EHTs (APD_30_: 254.6 ± 20.3 ms; APD_50_: 310.1 ± 22.8 ms; APD_80_: 342.3 ± 24.2 ms) and the IC-R406 EHTs (APD_30_: 193.8 ± 16.8 ms; APD_50_: 243.8 ± 32.7 ms; APD_80_: 282.9 ± 43.6 ms) (Supplementary Table 2). Taken together, the EHTs containing the *DES*^R406W^ variant displayed prolonged APDs.

### Transcriptomic and proteomic investigations of DES^R406W^ mutation consequences

To investigate how the *DES*^R406W^ mutation alters cell biology, transcriptomic and proteomic data were generated from 5 to 8 independent EHTs for all four hiPSC lines. Principal component analysis (PCA) highlighted that the principal factor in gene/protein expression variations, represented by PC1 (about 23% of gene and protein expression variation), mainly reflected differences between the 2 original hiPSCs lines (Control and PT-R406W; Figure 4A and B). The PC2, on the other hand, represented gene expression variations linked to the presence and absence of the *DES*^R406W^ mutation (Figure 4A and B).

**Figure 4.**
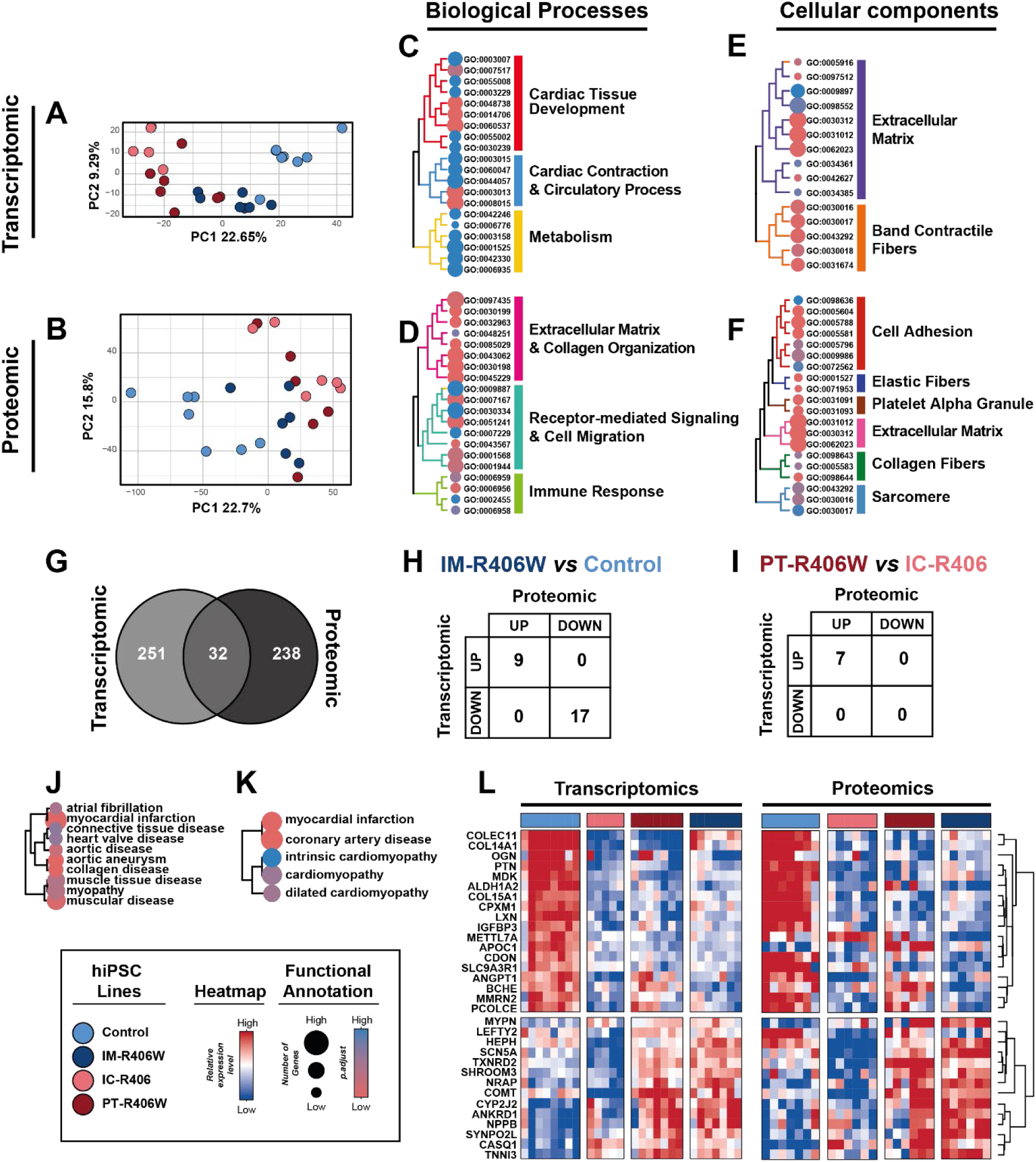
Transcriptomic and proteomic analyses point towards contractile and structural defects in *DES*^R406W^-EHTs. (Legend is found on the bottom left corner of the figure for definition of symbol color and size.) **A, B**. Principal Component Analysis (PCA) plot of transcriptomic and proteomic data from Control, IM-R406W, IC-R406W and PT-R406W EHTs. The percent of total variance due to the first two PCs is labeled. Five to eight EHTs were studied for each of the four hiPSC lines. **C-F**. Functional enrichment analysis of mRNAs (**C, E**) or proteins (**D, F**) with altered expression in IM-R406W compared to the Control EHTs and in PT-R406W compared to the IC-406W EHTs using the Gene Ontology categories “Biological Processes” (**C, D**) and “Cellular Components” (**E**, **F**). **G**. Shared dysregulated genes identified in transcriptomic and proteomic analyses, with pairwise comparisons shown in (**H**) and (**I**). Disease ontology annotation for (**J**) transcriptomic and (**K**) proteomic data. **L**. Heatmap showing the 32 dysregulated genes identified in both transcriptomic and proteomic analyses.

The expression of 210 genes and 200 proteins was significantly altered (≥ 2-fold change, with p-adjust≤ 0.01 and p-adjust≤ 0.05 for transcriptomics and proteomics, respectively) in IM-R406W EHTs *versus* Control, while 84 genes and 78 proteins showed an altered expression in PT-R406W *versus* IC-R406 (Supplementary Figure 1) among a total of 16846 genes and 7372 proteins detected (Supplementary Table 3_data 1-4). GO enrichment analysis of the transcriptomic signature related to the *DES*^R406W^ mutation – calculated on the union of all differentially expressed genes/proteins across IM-R406W EHTs *versus* Control and PT-R406W *versus* IC-R406 comparisons – pointed cardiac development, cardiomyocyte structure, contractile function, and metabolism as main over-represented biological processes and cellular components (Figure 4C, E). As expected, the *DES*^R406W^ mutation led to altered gene expression associated with the Z-disc (GO:0030018), the I-band (GO:0031674), and more broadly with extracellular matrix remodeling (GO:0031012) (Figure 4 C, E; Supplementary table 3_data 5-6). Consistent with the transcriptomic GO signature, the proteomic GO signature revealed not only extracellular matrix remodeling (GO:0030198) but also marked alterations in cell adhesion (GO:0031589; GO:0098636) and cell migration (GO:0043542) (Figure 4 D, F; Supplementary table 3_data 7-8). Overall, the functions over-represented in dysregulated transcripts and proteins were both related to the extracellular matrix and to the structural and contractile organization of EHTs. Interestingly, when the two pairs of isogenic lines were compared (*i.e.* IM-R406W *versus* Control and PT-R406W *versus* IC-R406), only 11 transcripts and 8 proteins were found to be similarly dysregulated in both comparisons (Supplementary Figure 2). This suggests that the DES p.R406W mutation induces variable remodeling depending on the genetic background of the hiPSC line. This observation is consistent with the literature, that the same mutation can lead a variety of phenotypes depending on the patients’ genetic background (Geryk and Charpentier 2024).

To further explore molecular alterations associated with the *DES*^R406W^ mutation, we performed an additional analysis on pooled -omics data from EHTs carrying the mutation (PT-R406W and IM-R406W) *versus* pooled data from those that did not (Control and IC-R406). We found that a total of 521 genes had their RNA and/or protein expression dysregulated, with 32 (6.14%) being concordantly altered in both transcriptomic and proteomic analyses (Figure 4G-I; Supplementary Figure 1 and Supplementary Table 3). A subset of these 521 transcripts/proteins has been associated with a large number of cardiac diseases (Figure 4J,K; Supplementary Table 3_data 9-10). The expression of the 32 genes significantly deregulated in both transcriptomic and proteomic datasets is displayed as a heatmap (Figure 4L). Interestingly, we observed a consistent upregulated expression of 14 genes across the *DES*^R406W^ mutant lines (lower part of the heatmap), independently of genetic background, including the cardiac sodium channel Na_V_1.5 (*SCN5A*), natriuretic peptide B (*NPPB*), and troponin I3 (*TNNI3*).

### Transmission electron microscopy

Desmin mutations often lead to structural changes of cardiomyocytes. To gain insight as to how the *DES*^R406W^ mutation affects the intracellular structure of EHTs, we imaged the tissues using transmission electron microscopy (TEM). We observed sarcomeres with prominent contractile filaments aligned at the Z-discs in both Control and IC-R406 EHTs (Figure 5, top panels) where the Z-discs were visible and distinguishable (black asterisks). For the EHTs carrying the *DES*^R406W^ mutation, PT-R406W and IM-R406W, although the sarcomeres were visible, the tissues exhibited Z-disc that were wider and “fuzzy” in appearance, making them almost indistinguishable from the filaments (black asterisks). (Figure 5, bottom panels). These results align with previous studies that used *DES* mutated mice (Herrmann et al. 2020).

**Figure 5.**
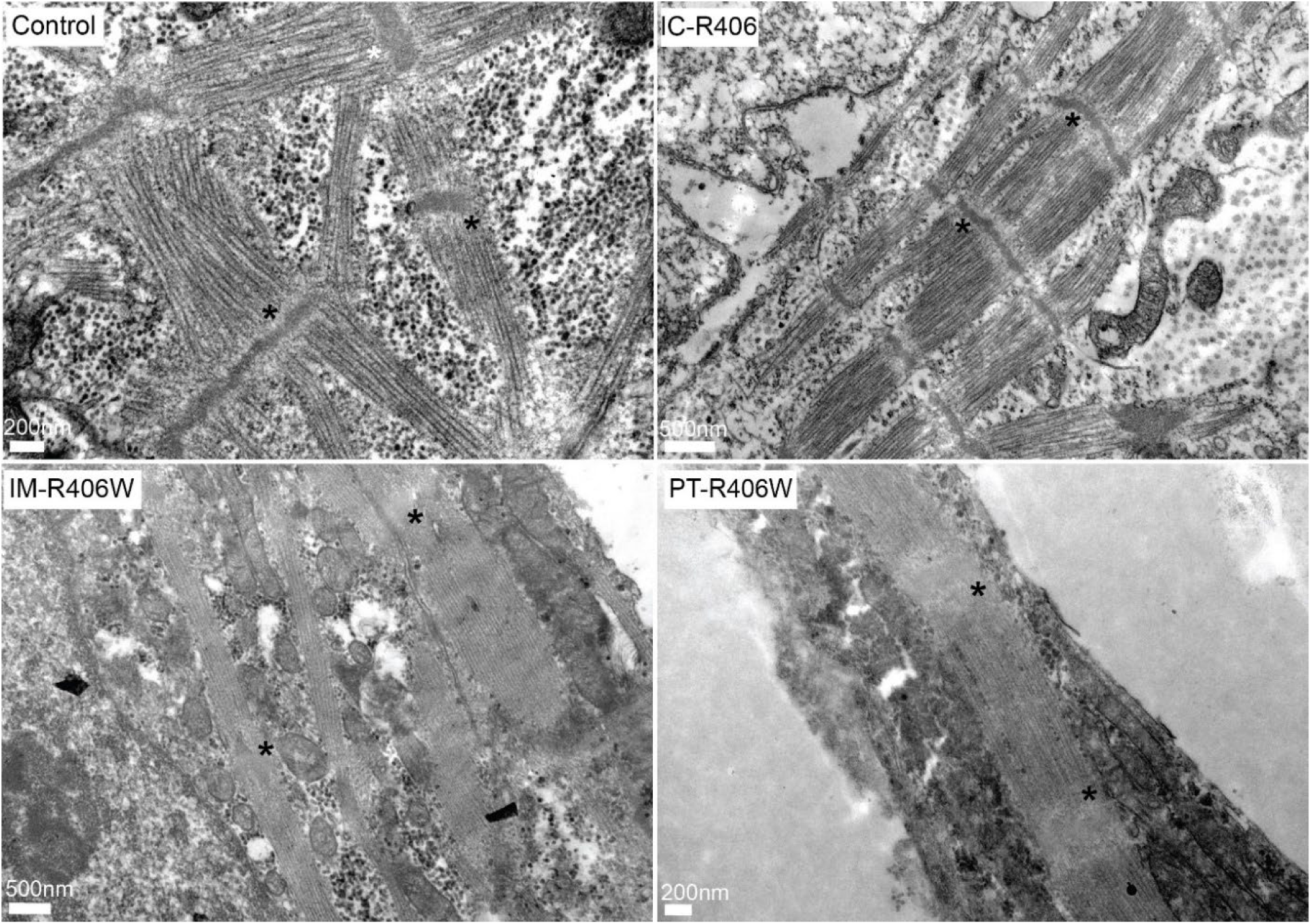
Transmission electron microscopy images of EHTs illustrate Z-disc abnormalities. The two control clones (Control and IC-R406; top panels) have visible Z-discs and filaments. The two clones carrying the *DES*^R406W^ variant (IM-R406W and PT-R406W; bottom panels) have wider Z-discs with a fuzzy appearance. The asterisks (black) highlight the Z-disc location and position.

As shown in Supplementary Figure 3, compared to the two control lines (Control and IC-R406), the PT-R406W and IM-R406W lines were characterized by a less striated desmin pattern and the desmoplakin signal is not as concentrated at the intercalated discs and is more dispersed. This shows that the *DES*^R406W^ mutation also altered intercalated disc structure of iPSC-CMs.

### Analysis of the cardiac function of *Des*^R405W^ knock-in (KI) mouse model

#### KI mice have increased susceptibility to ventricular tachyarrhythmias

To elucidate the consequences of the *DES*^R406W^ mutation on the incidence of cardiac arrhythmias, we investigated the cardiac electrical activity of a heterozygous KI mouse model carrying the equivalent p.R405W mutation. As shown in Table 1, KI mice were characterized by a moderately but significantly longer QRS complex, and lower R wave amplitude, than wildtype (WT) mice at both 10 and 20 weeks of age, suggesting that *Des* p.R405W mutation affects ventricular conduction. Similar results were observed in both males and females (Supplementary Table 4). No cardiac arrhythmia was observed during ECG recording in anaesthetized mice at both ages. However, as shown in Figure 6, ventricular tachyarrhythmias were more easily triggered with programmed electrical pacing in isolated hearts from 21-week-old KI mice than WT mice, especially under β-adrenergic stimulation. Not only was the incidence of arrhythmias higher in KI mice, but so was their severity, as only KI mice exhibited sustained ventricular tachycardia (VT). Arrhythmias were more easily and reproducibly triggered by S1S2 protocol, suggesting that they depended on reentrant mechanisms. To support this statement, we observed that cardiac impulse wavelengths, *i.e.,* the product of conduction velocity by refractory period, was shorter in KI mice compared to WT mice (Figure 6C).

**Table 1.** ECG parameters in 10- and 20-week-old wildtype (WT) and *Des*^R405W^ knock-in (KI) mice under baseline conditions.

|  | RR interval<br>(ms) | P wave<br>duration (ms) | PR interval<br>(ms) | QRS duration<br>(ms) | QT interval<br>(ms) | R wave<br>ampl. (mV) | S wave<br>ampl. (mV) |
| --- | --- | --- | --- | --- | --- | --- | --- |
| *10 weeks old* |  |  |  |  |  |  |  |
| **WT (n = 40)** | 123.4 ± 11.1 | 13.0 ± 1.8 | 38.3 ± 3.1 | 10.9 ± 0.9 | 53.2 ± 4.9 | 1.01 ± 0.23 | -0.3 ± 0.14 |
| **KI (n = 39)** | 126.7 ± 11.7 | 13.7 ± 1.7 | 38.5 ± 2.7 | 11.6 ± 1.2 ** | 54.4 ± 4.3 | 0.85 ± 0.25 ** | -0.2 ± 0.14 |
| *20 weeks old* |  |  |  |  |  |  |  |
| **WT (n = 25)** | 125.5 ± 11.7 | 13.0 ± 2.1 | 38.6 ± 2.9 | 10.5 ± 0.9 | 52.1 ± 3.3 | 0.84 ± 0.20 | -0.20 ± 0.12 |
| **KI (n = 19)** | 129.5 ± 14.7 | 12.7 ± 1.8 | 39.5 ± 2.1 | 11.6 ± 1.4 ** | 53.2 ± 3.9 | 0.71 ± 0.17 * | -0.29 ± 0.13 |

Abbreviation: ampl., amplitude. Data are expressed as mean ± standard deviation. \*, \*\*: p< 0.05 and p< 0.01, respectively, *versus* WT at corresponding age (Mann-Whitney test).

**Figure 6.**
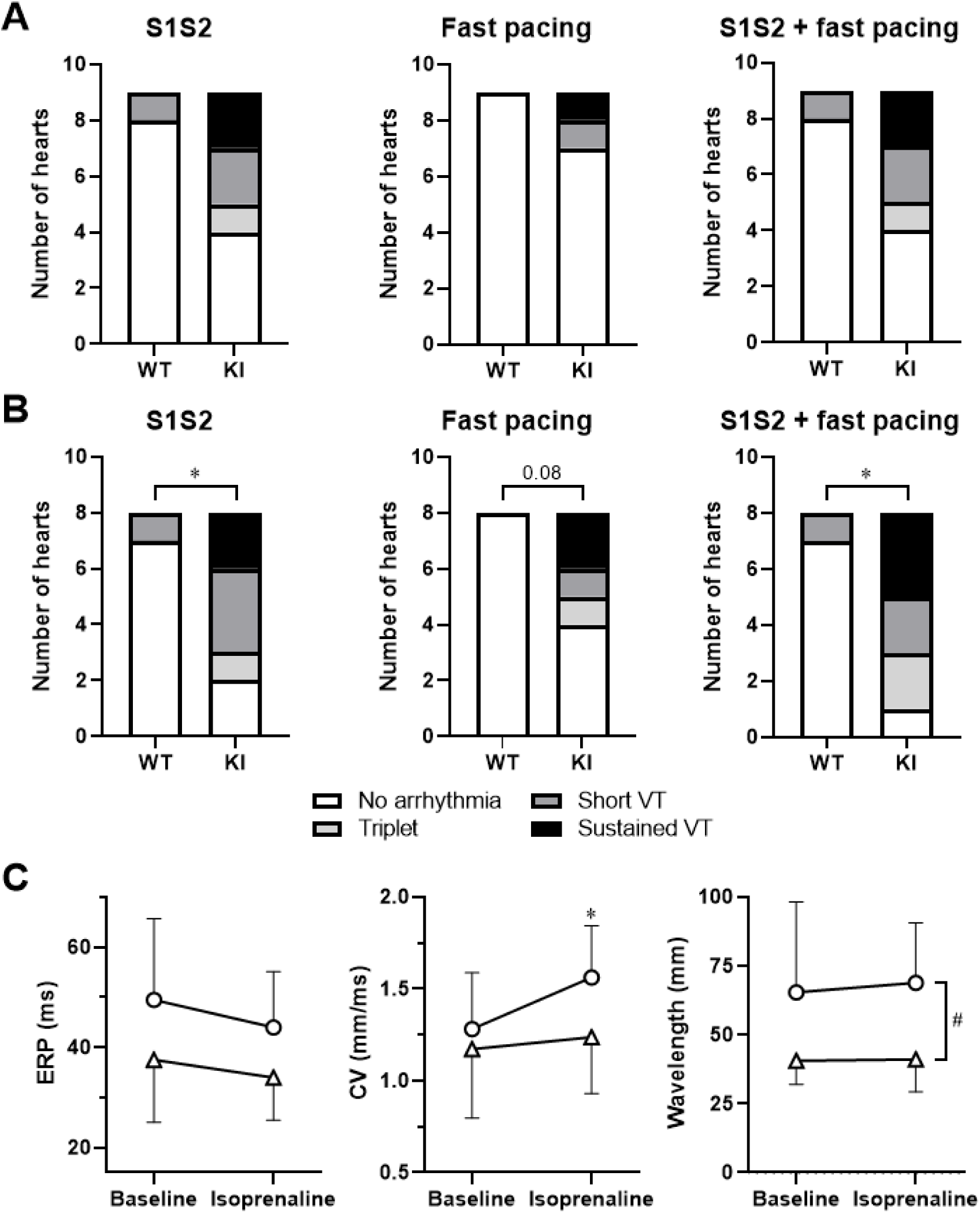
Pacing of isolated hearts triggered arrhythmia. **A, B.** Incidence of pacing-induced ventricular tachyarrhythmias in isolated hearts from wildtype (WT) and *Des*^R405W^ knock-in (KI) mice under baseline condition (**A**) and in the presence of 10^-7^ M isoprenaline (**B**). For each heart, the most severe arrhythmia was considered. See methods section for detailed description of the two pacing protocols. VT, ventricular tachycardia. *, p< 0.05 (Fisher’s exact test comparing “no arrhythmia” and “triplet+short VT+sustained VT” conditions). **C**.Ventricular effective refractory period (ERP), conduction velocity (CV) and wavelength at a pacing cycle length of 100 ms in isolated hearts from WT (circles; n = 8) and KI (triangles; n = 8) mice under baseline condition and in the presence of 10^-7^ M isoprenaline. *, p< 0.05 vs. baseline; #, p< 0.05 vs WT (two-way ANOVA completed by Sidak’s multiple comparisons test).

#### Female KI mice exhibit moderate cardiac hypertrophy

Table 2 shows the main echocardiographic parameters measured in 11- and 21-week-old WT and KI mice. Males and females were separated because they have different heart size. While there was no significant difference between WT and KI male mice, KI female mice showed signs of mild hypertrophy with a small, but significant decrease of the LV chamber diameter and a tendency for thicker LV wall in both diastole and systole, compared to WT mice. Similarly, the heart weight to tibia length ratio was significantly higher in 21-week-old KI female mice than in WT counterparts (Supplementary Table 5). Histological experiments performed at 21 weeks showed no statistical difference in fibrosis extent between WT and KI mice (Supplemental Table 6).

**Table 2.** Echocardiographic parameters in 11- and 21-week-old wildtype (WT) and *Des*^R405W^ knock-in (KI) mice under baseline conditions.

| Age | 11 weeks |  |  |  | 21 weeks |  |  |  |
| --- | --- | --- | --- | --- | --- | --- | --- | --- |
| Sex | Female |  | Male |  | Female |  | Male |  |
| Genotype (n) | WT (7) | KI (7) | WT (7) | KI (7) | WT (7) | KI (7) | WT (7) | KI (7) |
| **HR (bpm)** | 436 ± 58 | 421 ± 33 | 443 ± 55 | 445 ± 60 | 440 ± 51 | 449 ± 45 | 446 ± 58 | 451 ± 25 |
| **LVEF (%)** | 49.0 ± 8.3 | 54.5 ± 7.9 | 40.7 ± 7.4 | 41.9 ± 7.6 | 44.6 ± 6.5 | 51 ± 10 | 47.4 ± 9.0 | 44 ± 6 |
| **LVIDs (mm)** | 3.17 ± 0.40 | 2.75 ± 0.24* | 3.71 ± 0.38 | 3.57 ± 0.41 | 3.38 ± 0.42 | 2.90 ± 0.33* | 3.53 ± 0.53 | 3.51 ± 0.19 |
| **LVIDd (mm)** | 4.20 ± 0.26 | 3.82 ± 0.18** | 4.63 ± 0.32 | 4.48 ± 0.35 | 4.32 ± 0.41 | 3.91 ± 0.27* | 4.63 ± 0.44 | 4.48 ± 0.27 |
| **LVPWTs (mm)** | 0.91 ± 0.15 | 1.08 ± 0.17 | 0.81 ± 0.20 | 0.85 ± 0.08 | 0.92 ± 0.20 | 1.06 ± 0.20 | 0.92 ± 0.10 | 0.91 ± 0.09 |
| **LVPWTd (mm)** | 0.64 ± 0.10 | 0.78 ± 0.15* | 0.59 ± 0.15 | 0.65 ± 0.06 | 0.67 ± 0.16 | 0.78 ± 0.09 | 0.63 ± 0.10 | 0.66 ± 0.11 |

Abbreviations: HR, heart rate; LVEF, left ventricular ejection fraction; LVIDs, systolic left ventricular internal diameter; LVIDd, diastolic left ventricular internal diameter; LVPWTs, systolic left ventricular posterior wall thickness; LVPWTd, diastolic left ventricular posterior wall thickness. Data are expressed as mean ± standard deviation. \*, \*\*: p< 0.05 and p< 0.01, respectively, *versus* WT at corresponding age (Mann- Whitney test).

## Discussion

In this study, we identified the *DES*^R406W^ mutation in a young patient with occurrence of severe ventricular arrhythmias and SCD during her first decade of life. This variant (1) induces electrophysiological and structural abnormalities in hiPSC-CMs carrying the mutation; (2) leads to significant remodeling of the expression of genes and proteins involved in cardiac electrical activity and contractility, as well as its structural components; and (3) causes electrophysiological changes in mice carrying the equivalent *Des*^R405W^ mutation that increase arrhythmogenic risk, in the absence of fibrotic remodeling. More generally, this study highlights the value of the hiPSC-CMs and EHT models for analyzing the cellular and subcellular consequences in the heart, of mutations leading to desminopathies.

The *DES*^R406W^ mutation has been described earlier as a *de novo* mutation, *i.e.*, not found in index cases’ parents (Figure 1B) and other relatives (Dagvadorj et al. 2004), associated with severe cardiac and skeletal myopathy – although there are some exceptions (Arbustini et al. 2006) – which manifest early in life. One other case study has reported this variant in an 8-year-old female patient presenting with HCM and atrioventricular conduction block (Oka et al. 2021), highlighting the early onset and strong pathogenicity of this variant. Most *DES*^R406W^ carriers present with cardiac involvement first, followed by gradual skeletal muscle degeneration throughout life (Dagvadorj et al. 2004; Oka et al. 2021; Park et al. 2000). Patients are often diagnosed with RCM (Herrmann et al. 2020; Chen et al. 2021; Arbustini et al. 2006; Olivé et al. 2004), although HCM (Oka et al. 2021; Takegami et al. 2023) and DCM (Dagvadorj et al. 2004) have also been reported. The absence of an obvious structural cardiomyopathy in our patient might be linked to the early onset of her numerous VF episodes that ultimately led to early transplantation. Almost all patients with this mutation have cardiac conduction disorders (Herrmann et al. 2020; Olivé et al. 2004; Dagvadorj et al. 2004; Arbustini et al. 2006; Oka et al. 2021; Park et al. 2000; Chen et al. 2021), sometimes associated with ventricular tachyarrhythmias (Herrmann et al. 2020). Interestingly, the electrocardiographic pattern of the patient described in the present study resembles that observed in a recently identified inherited cardiac arrhythmia, the familial widespread ST-segment depression syndrome, which is characterized by persistent, non-ischemic ST-segment depression and increased risk of ventricular arrhythmias and SCD (Bundgaard et al. 2018; Christensen et al. 2021, 2022). The pathophysiological mechanisms underlying this syndrome have not been established and, to our knowledge, no causal gene has been identified thus far. A similar ECG pattern has also been found in a 15-year-old female patient with a homozygous deletion of 7 amino acids in the 1B helix domain of desmin (Piñol-Ripoll et al. 2009). A feature shared by these entities is the temporal stability of the ST-segment abnormality, in contrast to the dynamic ECGs of the Brugada and long-QT syndromes. A KI mouse model carrying the p.R405W mutation, the orthologous counterpart to the human p.R406W mutation, has been generated (Herrmann et al. 2020) but no electrophysiological investigations had been published, until now. This mouse, however, exhibits abnormal intercalated discs organization (Herrmann et al. 2020) that might partly explain the conduction defects and arrhythmias observed in patients carrying the *DES*^R406W^ mutation. Our study actually shows that these mice are more prone to develop ventricular arrhythmias than WT mice, even in the absence of severe cardiac structural remodeling.

Our study is the first to highlight the cardiac cellular electrophysiological consequences of the *DES*^R406W^ variant. Our results in human iPSC-CMs suggest that the mutation affects the repolarization process, which could contribute to the occurrence of arrhythmias. Surprisingly, we did not observe, in this model, any electrophysiological abnormalities that could explain the conduction defects observed in mice. For example, no change in AP upstroke velocity was observed. However, on the basis of previous studies of this variant showing desmosome alterations (Herrmann et al. 2020), which is confirmed here in micropatterned hiPSC-CMs, it would be useful to carry out experiments more specifically designed to measure the speed of electrical impulse conduction in EHTs.

Our multi-omics study of EHTs is one of the first to investigate the pathogenic mechanisms of desminopathies, especially the *DES*^R406W^ mutation. It provides a global overview of the altered functions in the cells. The transcriptomic data does not always represent the protein expression profiles, especially in pathology (Batoumeni et al. 2025; Haider and Pal 2013; Du et al. 2019; Liu et al. 2016). Therefore, additionally studying protein expression provides a more accurate view of the functions altered in desminopathy. However, with current technologies, transcriptomic sequencing technology provides broader coverage of the transcriptome than proteomic approaches can achieve for the proteome (Haider and Pal 2013). This coverage difference is observed in the present study, with close to 8 000 proteins compared to almost 15 000 genes. Therefore, we decided to use both techniques. The transcriptomic and proteomic data generated in this study suggest that the hiPSC-CM-derived EHT model carrying the *DES*^R406W^ mutation recapitulates a number of features characteristic of desminopathies, with biological processes and cellular components associated to structural remodeling of cardiomyocytes, particularly through disruption of the contractile apparatus. Concomitant alterations in extracellular matrix, including collagen disorganization, and defective cell-to-cell connections further indicate compromised tissue integrity that may result in tissue replacement. Accordingly, analysis of TEM images revealed a faint appearance of the Z-discs in the mutant tissues. This is intriguing since the biological process GO terms revealed an upregulation of proteins related to cardiac contraction, such as Troponin I, Myozenin-1 and Myopalladin (Supplementary Tables 3 and 8). This suggests a positive feedback upregulation that cannot counteract the effect of the *DES*^R406W^ mutation. Decreased levels of desmin at the Z-discs have been reported in desmin KI R405W mice (Herrmann et al. 2020), supporting the notion that the interplay between desmin and its cell-spanning connections is critical for proper cardiac function. Accordingly, the multi-omics study also shows that changes in the structure of desmin filaments lead not only to structural alterations in the cell but also to a complex remodeling of the expression of a large number of transcripts and proteins implicated in many cardiomyocyte functions. From a structure-function relationship aspect, we saw DEGs related to cardiac muscle tissue development (GO:0048738) that highlighted changes in key genes important for cardiac function such as *SCN5A, GJA1, MYL2,* and *MYH7.* The desmin gene was also found to be differentially expressed along with *TNNI3* (GO: 0007517), tying them together in the context of cell function and integrity and highlighting the pathogenicity of the mutation.

Interestingly, the genetic backgrounds of the hiPSC lines appeared to influence the outcomes of this study. As mentioned above, a single desmin mutation can lead to different cardiomyopathies, with varying severity and age of onset (Geryk and Charpentier 2024). Importantly, our results demonstrated differences in gene remodeling depending on the origin of the hiPSC line (patient-derived *versus* control-derived cells). Although similar trends were observed and divide the EHTs based on the presence or absence of the *DES*^R406W^ mutation, the genetic background clearly influenced the extent and nature of the molecular remodeling. Such variability may contribute to the development of distinct cardiomyopathic phenotypes. Altogether, these findings highlight the importance of using isogenic hiPSC lines to perform reliable and interpretable analyses.

Here, we demonstrated that EHTs could be a relevant model for studying the cardiac consequences of desminopathy. Our study illustrates that hiPSC-CMs are a relevant model to investigate desmin mutations. Furthermore, EHTs generated from these CMs are a valuable three-dimensional model that can recapitulate certain aspects of the disease. To date, studies that have investigated desmin used different cell lines that are typically desmin- and vimentin-free, such as the human adrenocortical carcinoma (SW13) cells (Goudeau et al. 2006; Bär et al. 2005; Chourbagi et al. 2011). These studies provided insight into the structural consequences of mutant desmin filaments. It was established that the *DES*^R406W^ variant forms aggregates and disrupts proper filament formation (Goudeau et al. 2006; Bär et al. 2005). The aggregate phenotype has also been observed in biopsy samples from patients with the *DES*^R406W^ mutation (Arbustini et al. 2006; Park et al. 2000; Takegami et al. 2023; Olivé et al. 2004). Nevertheless, transfected hiPSC-CMs overexpressing the *DES*^R406W^ variant have demonstrated aggregate deposits in these cells (Kubánek et al. 2020), providing support for modeling desminopathy with hiPSC-CMs. Mutations located in different parts of *DES* have likewise been transfected into hiPSC-CMs, where aggregates were observed (Kulikova et al. 2021; Protonotarios et al. 2021; Brodehl et al. 2019), demonstrating pathogenicity of the variants. However, few studies have used patient hiPSCs with native *DES* expression level, to investigate the molecular mechanisms of desminopathies (Hovhannisyan et al. 2024; Tse et al. 2013; Batoumeni et al. 2025). One limitation of hiPSC-CMs is their immaturity since they have a metabolic, structural and electrophysiological phenotype that differs from that of adult cardiomyocytes (Ronaldson-Bouchard et al. 2018; Karbassi et al. 2020; Bekhite and Schulze 2021; Wu et al. 2021). On the other hand, this limitation might represent an advantage if one is interested in the developmental consequences of gene mutations leading to early-onset diseases, such as the *DES*^R406W^ mutation. From an electrophysiological point of view, for the purpose of this study we have been able to recapitulate certain aspects of mature cardiac AP with the dynamic clamp technique in order to compare pertinent electrophysiological parameters. We also found that the electrical maturity of EHTs could allow better identification of electrophysiological abnormalities, although our study would require more in-depth exploitation of this model.

Finally, our electrophysiological experiments on the *Des*^R405W^ mice demonstrate, for the first time, that the mutation affects ventricular conduction and increases susceptibility to ventricular tachyarrhythmias in the absence of fibrotic remodeling, although incipient hypertrophy was observed. These results are consistent with the patient’s phenotype. The repolarization defects observed in hiPSC models were not found in the mouse model. However, this is not surprising, as the ion currents involved in cardiac repolarization in mice differ significantly from those responsible for repolarization in humans (Nerbonne and Kass 2005).

In conclusion, this study demonstrates that the desmin p.R406W variant is a highly pathogenic mutation responsible for severe ventricular arrhythmias and sudden cardiac death from an early age. The integration of human cell models and a mouse model has shown that the mutation causes a global dysregulation of genes and proteins involved in cardiac function and cell adhesion, induces specific electrophysiological defects, and increases vulnerability to arrhythmias. This work highlights the relevance of hiPSC-derived cardiomyocytes and 3D EHT models for modeling human desminopathies and differentiating their sub-phenotypes.

## Acknowledgements

We are most grateful to the *Centre National de Recherche en Génomique Humaine, Institut de Génomique, CEA, Evry, France,* for whole genome sequencing, and to the Genomics Core Facility GenoA, member of Biogenouest and *France Génomique* and to the Bioinformatics Core Facility BiRD, member of Biogenouest and *Institut Français de Bioinformatique* (ANR-11-INBS-0013) for the use of their resources and their technical support. We would like to thank Rodolphe Perrot from SCIAM (Common Service for Imaging and Microscopy Analysis, University of Angers, France) for TEM sample preparation and observation. The authors acknowledge THERASSAY core facility (SFR Bonamy, Nantes, France), the Prot’ICO proteomics facility (Institut de Cancérologie de l’Ouest, Angers, France), both members of the Scientific Interest Group (GIS) Biogenouest and IBISA, for the use of the echograph and technical support.

## References

1. Agnetti, Giulio, Harald Herrmann, and Shenhav Cohen. 2022. “New Roles for Desmin in the Maintenance of Muscle Homeostasis.” The FEBS Journal 289 (10): 2755–70. 10.1111/febs.15864.

2. Al Sayed, Zeina R., Robin Canac, Bastien Cimarosti, et al. 2021. “Human Model of *IRX5* Mutations Reveals Key Role for This Transcription Factor in Ventricular Conduction.” Cardiovascular Research 117 (9): 2092–107. 10.1093/cvr/cvaa259.

3. Al Sayed, Zeina R., Mariam Jouni, Jean-Baptiste Gourraud, et al. 2021. “A Consistent Arrhythmogenic Trait in Brugada Syndrome Cellular Phenotype.” Clinical and Translational Medicine 11 (6): e413. 10.1002/ctm2.413.

4. Arbustini, Eloisa, Michele Pasotti, Andrea Pilotto, et al. 2006. “Desmin Accumulation Restrictive Cardiomyopathy and Atrioventricular Block Associated with Desmin Gene Defects.” European Journal of Heart Failure 8 (5): 477–83. 10.1016/j.ejheart.2005.11.003.

5. Bär, H., N. Mucke, A. Kostareva, G. Sjoberg, U. Aebi, and H. Herrmann. 2005. “Severe Muscle Disease-Causing Desmin Mutations Interfere with in Vitro Filament Assembly at Distinct Stages.” Proceedings of the National Academy of Sciences 102 (42): 15099–104. 10.1073/pnas.0504568102.

6. Batoumeni, Vivien, Yeranuhi Hovhannisyan, Bénédicte Gobert, et al. 2025. “Integrated Phenotypic and Transcriptomic Characterization of Desmin-Related Cardiomyopathy in hiPSC-Derived Cardiomyocytes and Machine Learning-Based Classification of Disease Features.” European Journal of Cell Biology 104 (3): 151502. 10.1016/j.ejcb.2025.151502.

7. Bekhite, Mohamed M., and P. Christian Schulze. 2021. “Human Induced Pluripotent Stem Cell as a Disease Modeling and Drug Development Platform—A Cardiac Perspective.” Cells 10 (12): 12. 10.3390/cells10123483.

8. Bermúdez-Jiménez, Francisco José, Víctor Carriel, Andreas Brodehl, et al. 2018. “Novel Desmin Mutation p.Glu401Asp Impairs Filament Formation, Disrupts Cell Membrane Integrity, and Causes Severe Arrhythmogenic Left Ventricular Cardiomyopathy/Dysplasia.” Circulation 137 (15): 1595–610. 10.1161/CIRCULATIONAHA.117.028719.

9. Blighe, Kevin, Jared Andrews, Sharmila Rana, et al. 2018. EnhancedVolcano. Bioconductor, released. 10.18129/B9.BIOC.ENHANCEDVOLCANO.

10. Boukens, Bastiaan J., Mathilde R. Rivaud, Stacey Rentschler, and Ruben Coronel. 2014. “Misinterpretation of the Mouse ECG: ‘Musing the Waves of Mus Musculus.’” The Journal of Physiology 592 (Pt 21): 4613–26. 10.1113/jphysiol.2014.279380.

11. Brodehl, Andreas, Seyed Ahmad Pour Hakimi, Caroline Stanasiuk, et al. 2019. “Restrictive Cardiomyopathy Is Caused by a Novel Homozygous Desmin (DES) Mutation p.Y122H Leading to a Severe Filament Assembly Defect.” Genes 10 (11): 918. 10.3390/genes10110918.

12. Bundgaard, Henning, Christian Jøns, Elisabeth M. Lodder, et al. 2018. “A Novel Familial Cardiac Arrhythmia Syndrome with Widespread ST-Segment Depression.” New England Journal of Medicine 379 (18): 1780–81. 10.1056/NEJMc1807668.

13. Calloe, Kirstine, Michelle Geryk, Kristine Freude, et al. 2022. “The G213D Variant in Nav1.5 Alters Sodium Current and Causes an Arrhythmogenic Phenotype Resulting in a Multifocal Ectopic Purkinje-Related Premature Contraction Phenotype in Human-Induced Pluripotent Stem Cell-Derived Cardiomyocytes.” EP Europace 24 (12): 2015–27. 10.1093/europace/euac090.

14. Capetanaki, Yassemi, Robert J. Bloch, Asimina Kouloumenta, Manolis Mavroidis, and Stelios Psarras. 2007. “Muscle Intermediate Filaments and Their Links to Membranes and Membranous Organelles.” *Experimental Cell Research*, Special Issue - Intermediate Filaments, vol. 313 (10): 2063–76. 10.1016/j.yexcr.2007.03.033.

15. Charpentier, Eric, Marine Cornec, Solenne Dumont, et al. 2021. *3’ RNA Sequencing for Robust and Low-Cost Gene Expression Profiling*. Preprint. Protocol Exchange. 10.21203/rs.3.pex-1336/v1.

16. Chen, Zixian, Rui Li, Yongxiang Wang, et al. 2021. “Features of Myocardial Injury Detected by Cardiac Magnetic Resonance in a Patient with Desmin-related Restrictive Cardiomyopathy.” ESC Heart Failure 8 (6): 5560–64. 10.1002/ehf2.13624.

17. Chourbagi, Oussama, Francine Bruston, Marianna Carinci, et al. 2011. “Desmin Mutations in the Terminal Consensus Motif Prevent Synemin-Desmin Heteropolymer Filament Assembly.” Experimental Cell Research 317 (6): 886–97. 10.1016/j.yexcr.2011.01.013.

18. Christensen, Alex Hørby, Benjamin Chris Nyholm, Christoffer Rasmus Vissing, et al. 2021. “Natural History and Clinical Characteristics of the First 10 Danish Families With Familial ST-Depression Syndrome.” Journal of the American College of Cardiology 77 (20): 2617–19. 10.1016/j.jacc.2021.03.313.

19. Christensen, Alex Hørby, Christoffer Rasmus Vissing, Adrian Pietersen, et al. 2022. “Electrocardiographic Findings, Arrhythmias, and Left Ventricular Involvement in Familial ST-Depression Syndrome.” Circulation: Arrhythmia and Electrophysiology 15 (4): e010688. 10.1161/CIRCEP.121.010688.

20. Dagvadorj, Ayush, Montse Olivé, Jean-Andoni Urtizberea, et al. 2004. “A Series of West European Patients with Severe Cardiac and Skeletal Myopathy Associated with a de Novo R406W Mutation in Desmin.” Journal of Neurology 251 (2): 143–49. 10.1007/s00415-004-0289-3.

21. Derangeon, Mickael, Jérôme Montnach, Cynthia Ore Cerpa, et al. 2017. “Transforming Growth Factor β Receptor Inhibition Prevents Ventricular Fibrosis in a Mouse Model of Progressive Cardiac Conduction Disease.” Cardiovascular Research 113 (5): 464– 74. 10.1093/cvr/cvx026.

22. Du, Yina, Geremy C. Clair, Denise Al Alam, et al. 2019. “Integration of Transcriptomic and Proteomic Data Identifies Biological Functions in Cell Populations from Human Infant Lung.” American Journal of Physiology. Lung Cellular and Molecular Physiology 317 (3): L347–60. 10.1152/ajplung.00475.2018.

23. Geryk, Michelle, Robin Canac, Virginie Forest, et al. 2024. “Generation of a Patient-Specific Induced Pluripotent Stem Cell Line Carrying the DES p.R406W Mutation, an Isogenic Control and a DES p.R406W Knock-in Line.” Stem Cell Research 77 (June): 103396. 10.1016/j.scr.2024.103396.

24. Geryk, Michelle, and Flavien Charpentier. 2024. “Pathophysiological Mechanisms of Cardiomyopathies Induced by Desmin Gene Variants Located in the C-Terminus of Segment 2B.” Journal of Cellular Physiology 239 (5): e31254. 10.1002/jcp.31254.

25. Girardeau, Aurore, Diane Atticus, Robin Canac, et al. 2022. “Generation of Human Induced Pluripotent Stem Cell Lines from Four Unrelated Healthy Control Donors Carrying European Genetic Background.” Stem Cell Research 59 (March): 102647. 10.1016/j.scr.2021.102647.

26. Goudeau, Bertrand, Fernando Rodrigues-Lima, Dirk Fischer, et al. 2006. “Variable Pathogenic Potentials of Mutations Located in the Desmin Alpha-Helical Domain.” Human Mutation 27 (9): 906–13. 10.1002/humu.20351.

27. Gu, Zuguang. 2022. “Complex Heatmap Visualization.” iMeta 1 (3): e43. 10.1002/imt2.43.

28. Haider, Saad, and Ranadip Pal. 2013. “Integrated Analysis of Transcriptomic and Proteomic Data.” Current Genomics 14 (2): 91–110. 10.2174/1389202911314020003.

29. Hakibilen, Coralie, Florence Delort, Marie-Thérèse Daher, et al. 2022. “Desmin Modulates Muscle Cell Adhesion and Migration.” Frontiers in Cell and Developmental Biology 10: 783724. 10.3389/fcell.2022.783724.

30. Herrmann, Harald, Eva Cabet, Nicolas R. Chevalier, et al. 2020. “Dual Functional States of R406W-Desmin Assembly Complexes Cause Cardiomyopathy With Severe Intercalated Disc Derangement in Humans and in Knock-In Mice.” Circulation 142 (22): 2155–71. 10.1161/CIRCULATIONAHA.120.050218.

31. Hol, Elly M., and Yassemi Capetanaki. 2017. “Type III Intermediate Filaments Desmin, Glial Fibrillary Acidic Protein (GFAP), Vimentin, and Peripherin.” Cold Spring Harbor Perspectives in Biology 9 (12): a021642. 10.1101/cshperspect.a021642.

32. Hovhannisyan, Yeranuhi, Zhenlin Li, Domitille Callon, et al. 2024. “Critical Contribution of Mitochondria in the Development of Cardiomyopathy Linked to Desmin Mutation.” Stem Cell Research & Therapy 15 (1): 10. 10.1186/s13287-023-03619-7

33. Howie, Bryan N., Peter Donnelly, and Jonathan Marchini. 2009. “A Flexible and Accurate Genotype Imputation Method for the Next Generation of Genome-Wide Association Studies.” PLOS Genetics 5 (6): e1000529. 10.1371/journal.pgen.1000529.

34. Karbassi, Elaheh, Aidan Fenix, Silvia Marchiano, et al. 2020. “Cardiomyocyte Maturation: Advances in Knowledge and Implications for Regenerative Medicine.” Nature Reviews Cardiology 17 (6): 6. 10.1038/s41569-019-0331-x.

35. Kubánek, Miloš, Tereza Schimerová, Lenka Piherová, et al. 2020. “Desminopathy: Novel Desmin Variants, a New Cardiac Phenotype, and Further Evidence for Secondary Mitochondrial Dysfunction.” Journal of Clinical Medicine 9 (4): 937. 10.3390/jcm9040937.

36. Kulikova, Olga, Andreas Brodehl, Anna Kiseleva, et al. 2021. “The Desmin (DES) Mutation p.A337P Is Associated with Left-Ventricular Non-Compaction Cardiomyopathy.” Genes 12 (1): 121. 10.3390/genes12010121.

37. Lam, Chi Keung, Lei Tian, Nadjet Belbachir, et al. 2019. “Identifying the Transcriptome Signatures of Calcium Channel Blockers in Human Induced Pluripotent Stem Cell-Derived Cardiomyocytes.” Circulation Research 125 (2): 212–22. 10.1161/CIRCRESAHA.118.314202.

38. Lê, Sébastien, Julie Josse, and François Husson. 2008. “FactoMineR: An R Package for Multivariate Analysis.” Journal of Statistical Software 25 (March): 1–18. 10.18637/jss.v025.i01.

39. Liu, Yansheng, Andreas Beyer, and Ruedi Aebersold. 2016. “On the Dependency of Cellular Protein Levels on mRNA Abundance.” Cell 165 (3): 535–50. 10.1016/j.cell.2016.03.014.

40. Love, Michael I., Wolfgang Huber, and Simon Anders. 2014. “Moderated Estimation of Fold Change and Dispersion for RNA-Seq Data with DESeq2.” Genome Biology 15 (12): 550. 10.1186/s13059-014-0550-8.

41. Mannhardt, Ingra, Umber Saleem, Anika Benzin, et al. 2017. “Automated Contraction Analysis of Human Engineered Heart Tissue for Cardiac Drug Safety Screening.” Journal of Visualized Experiments, no. 122 (April): 55461. 10.3791/55461.

42. Mavroidis, Manolis, Panagiota Panagopoulou, Ioanna Kostavasili, Noah Weisleder, and Yassemi Capetanaki. 2008. “A Missense Mutation in Desmin Tail Domain Linked to Human Dilated Cardiomyopathy Promotes Cleavage of the Head Domain and Abolishes Its Z-Disc Localization.” The FASEB Journal 22 (9): 3318–27. 10.1096/fj.07-088724.

43. Meijer van Putten, Rosalie M. E., Isabella Mengarelli, Kaomei Guan, et al. 2015. “Ion Channelopathies in Human Induced Pluripotent Stem Cell Derived Cardiomyocytes: A Dynamic Clamp Study with Virtual IK1.” Frontiers in Physiology 6 (February). 10.3389/fphys.2015.00007.

44. Milner, D. J., G. Weitzer, D. Tran, A. Bradley, and Y. Capetanaki. 1996. “Disruption of Muscle Architecture and Myocardial Degeneration in Mice Lacking Desmin.” The Journal of Cell Biology 134 (5): 1255–70. 10.1083/jcb.134.5.1255.

45. Nerbonne, Jeanne M., and Robert S. Kass. 2005. “Molecular Physiology of Cardiac Repolarization.” Physiological Reviews 85 (4): 1205–53. 10.1152/physrev.00002.2005.

46. Oka, Hideharu, Kouichi Nakau, Rina Imanishi, et al. 2021. “A Case Report of a Rare Heterozygous Variant in the Desmin Gene Associated With Hypertrophic Cardiomyopathy and Complete Atrioventricular Block.” CJC Open 3 (9): 1195–98. 10.1016/j.cjco.2021.05.003.

47. Olivé, Montse, Lev Goldfarb, Dolores Moreno, et al. 2004. “Desmin-Related Myopathy: Clinical, Electrophysiological, Radiological, Neuropathological and Genetic Studies.” Journal of the Neurological Sciences 219 (1): 125–37. 10.1016/j.jns.2004.01.007.

48. Park, Kye-Yoon, Marinos C. Dalakas, Christina Semino-Mora, et al. 2000. “Sporadic Cardiac and Skeletal Myopathy Caused by a de Novo Desmin Mutation.” Clinical Genetics 57 (6): 423–29. 10.1034/j.1399-0004.2000.570604.x.

49. Perez-Riverol, Yasset, Chakradhar Bandla, Deepti J. Kundu, et al. 2025. “The PRIDE Database at 20 Years: 2025 Update.” Nucleic Acids Research 53 (D1): D543–53. 10.1093/nar/gkae1011.

50. Piñol-Ripoll, Gerard, Alexey Shatunov, Ana Cabello, et al. 2009. “Severe Infantile-Onset Cardiomyopathy Associated with a Homozygous Deletion in Desmin.” Neuromuscular Disorders 19 (6): 418–22. 10.1016/j.nmd.2009.04.004.

51. Protonotarios, Alexandros, Andreas Brodehl, Angeliki Asimaki, et al. 2021. “The Novel Desmin Variant p.Leu115Ile Is Associated With a Unique Form of Biventricular Arrhythmogenic Cardiomyopathy.” Canadian Journal of Cardiology 37 (6): 857–66. 10.1016/j.cjca.2020.11.017.

52. Ritchie, Matthew E., Belinda Phipson, Di Wu, et al. 2015. “Limma Powers Differential Expression Analyses for RNA-Sequencing and Microarray Studies.” Nucleic Acids Research 43 (7): e47. 10.1093/nar/gkv007.

53. Ronaldson-Bouchard, Kacey, Stephen P. Ma, Keith Yeager, et al. 2018. “Advanced Maturation of Human Cardiac Tissue Grown from Pluripotent Stem Cells.” Nature 556 (7700): 7700. 10.1038/s41586-018-0016-3.

54. Ronaldson-Bouchard, Kacey, Keith Yeager, Diogo Teles, et al. 2019. “Engineering of Human Cardiac Muscle Electromechanically Matured to an Adult-like Phenotype.” Nature Protocols 14 (10): 2781–817. 10.1038/s41596-019-0189-8.

55. Royer, Anne, Toon A. B. van Veen, Sabrina Le Bouter, et al. 2005. “Mouse Model of SCN5A-Linked Hereditary Lenègre’s Disease.” Circulation 111 (14): 1738–46. 10.1161/01.CIR.0000160853.19867.61.

56. Schindelin, Johannes, Ignacio Arganda-Carreras, Erwin Frise, et al. 2012. “Fiji - an Open Source Platform for Biological Image Analysis.” Nature Methods 9 (7): 10.1038/nmeth.2019. 10.1038/nmeth.2019.

57. Smolina, Natalia, Joseph Bruton, Gunnar Sjoberg, Anna Kostareva, and Thomas Sejersen. 2014. “Aggregate-Prone Desmin Mutations Impair Mitochondrial Calcium Uptake in Primary Myotubes.” Cell Calcium 56 (4): 269–75. 10.1016/j.ceca.2014.08.001.

58. Smolina, Natalia, Aleksandr Khudiakov, Anastasiya Knyazeva, et al. 2020. “Desmin Mutations Result in Mitochondrial Dysfunction Regardless of Their Aggregation Properties.” Biochimica et Biophysica Acta (BBA) - Molecular Basis of Disease 1866 (6): 165745. 10.1016/j.bbadis.2020.165745.

59. Takegami, Naoki, Akihiko Mitsutake, Tatsuo Mano, et al. 2023. “The Myocardial Accumulation of Aggregated Desmin Protein in a Case of Desminopathy with a *de Novo DES* p.R406W Mutation.” Internal Medicine, 0992–22. 10.2169/internalmedicine.0992-22.

60. Treat, Jacqueline A., Robert J. Goodrow, Corina T. Bot, Rodolfo J. Haedo, and Jonathan M. Cordeiro. 2019. “Pharmacological Enhancement of Repolarization Reserve in Human Induced Pluripotent Stem Cells Derived Cardiomyocytes.” Biochemical Pharmacology 169 (November): 113608. 10.1016/j.bcp.2019.08.010.

61. Tse, Hung-Fat, Jenny C. Y. Ho, Shing-Wan Choi, et al. 2013. “Patient-Specific Induced-Pluripotent Stem Cells-Derived Cardiomyocytes Recapitulate the Pathogenic Phenotypes of Dilated Cardiomyopathy Due to a Novel DES Mutation Identified by Whole Exome Sequencing.” Human Molecular Genetics 22 (7): 1395–403. 10.1093/hmg/dds556.

62. Tsikitis, Mary, Zoi Galata, Manolis Mavroidis, Stelios Psarras, and Yassemi Capetanaki. 2018. “Intermediate Filaments in Cardiomyopathy.” Biophysical Reviews 10 (4): 1007–31. 10.1007/s12551-018-0443-2.

63. Wilders, Ronald. 2006. “Dynamic Clamp: A Powerful Tool in Cardiac Electrophysiology: Dynamic Clamp: A Powerful Tool in Cardiac Electrophysiology.” The Journal of Physiology 576 (2): 349–59. 10.1113/jphysiol.2006.115840.

64. Wu, Peng, Gang Deng, Xiyalatu Sai, Huiming Guo, Huanlei Huang, and Ping Zhu. 2021. “Maturation Strategies and Limitations of Induced Pluripotent Stem Cell-Derived Cardiomyocytes.” Bioscience Reports 41 (6): BSR20200833. 10.1042/BSR20200833.

65. Xu, Shuangbin, Erqiang Hu, Yantong Cai, et al. 2024. “Using clusterProfiler to Characterize Multiomics Data.” Nature Protocols 19 (11): 3292–320. 10.1038/s41596-024-01020-z.

