## Supplementary material for "Desmin p.R406W mutation is associated with arrhythmias through structural and electrophysiological remodeling": Data supplement

**Geryk M et al.**

### Supplementary Data

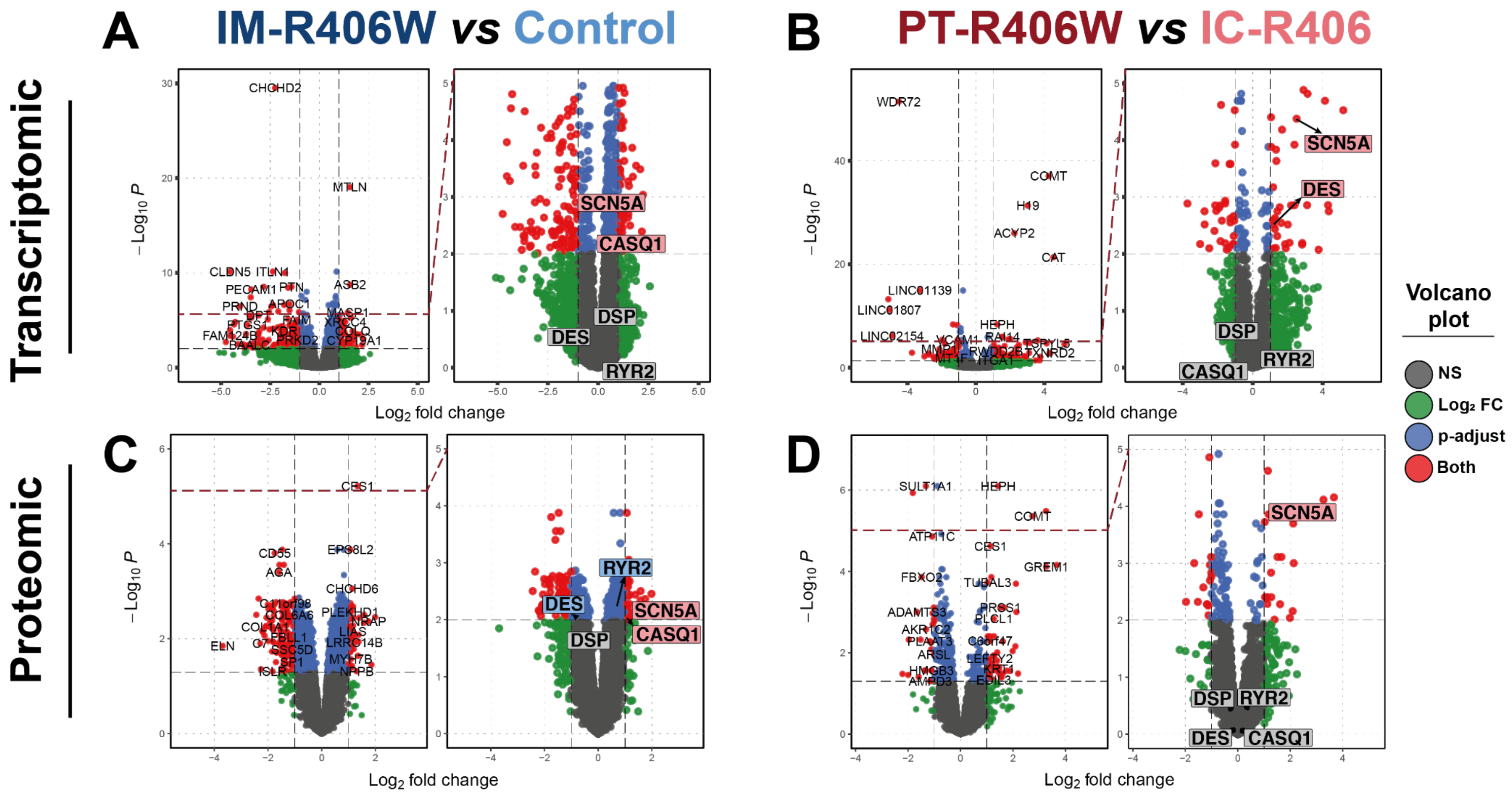


**Supplementary Figure 1.** Volcano plots representing the differentially expressed genes (A,B) or proteins (C,D) in IM-R406W versus Control EHTs (**A, C**) and in PT-R406W *versus* IC-406W EHTs (**B**,**D**). For each panel, the plots on the right are enlarged versions of the plots on the left. Grey dots (NS): non-significant changes with a log2 fold change < 1; blue dots (p-adjust): statistically significant changes with a log2 fold change < 1; green dots (Log2 FC): non-significant changes with a log2 fold change > 1; red dots (Both): statistically significant changes with a log2 fold change > 1.

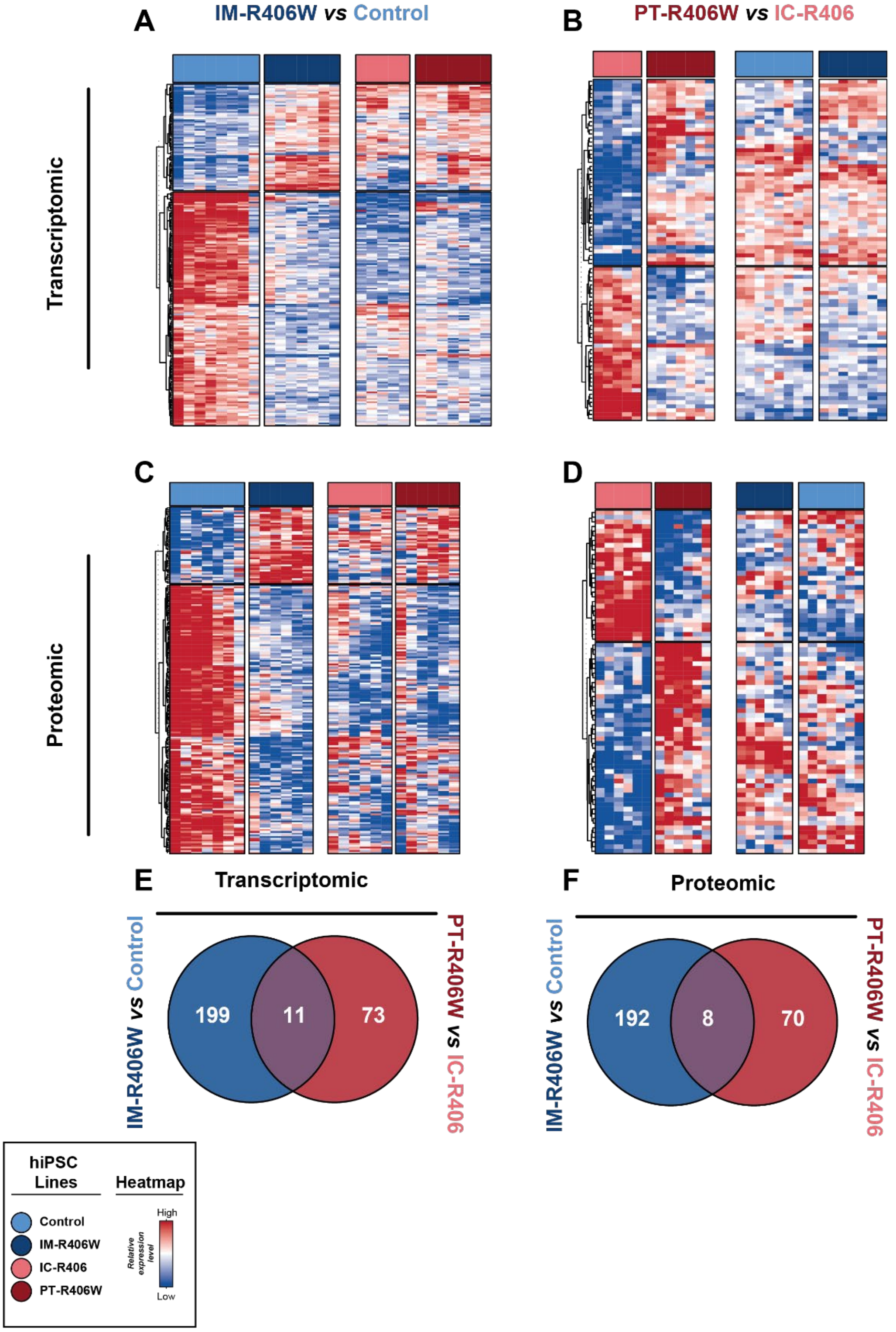


**Supplementary Figure 2. A-D.** Heatmaps showing the differentially expressed genes (A,B) or proteins (C,D) in IM-R406W *versus* Control EHTs (**A, C**) and in PT-R406W *versus* IC-406W EHTs (heatmaps **B**,**D**). In each heatmap, the first two columns correspond to the comparison from which the differentially expressed genes/proteins were identified, whereas the last two columns show the expression pattern of the same genes/proteins in the other p.R406W mutant line and its corresponding control for comparison. **E-F.** Venn diagrams showing the numbers of dysregulated genes (E) and proteins (F) in IM-R406W *versus* Control EHTs and PT-R406W *versus* IC-406 EHTs. Only 11 genes and 8 proteins are commonly dysregulated in the two p.R406W mutant lines.

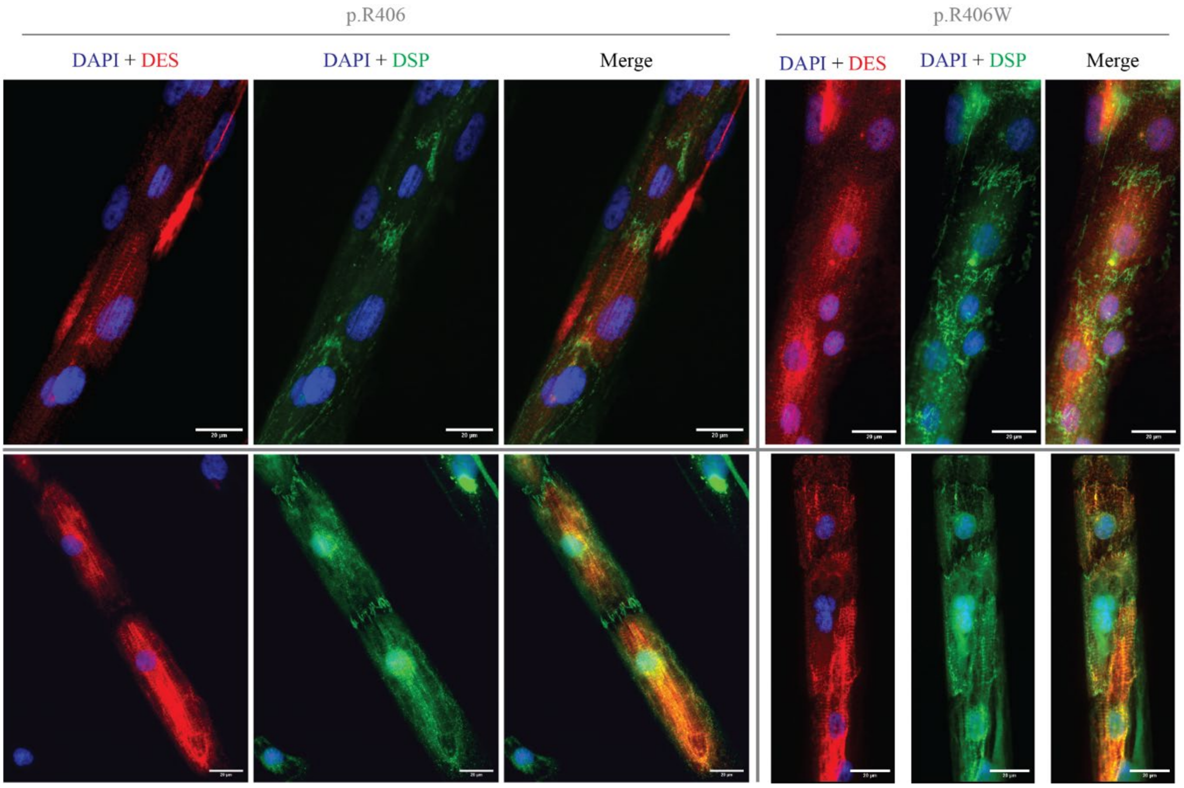


**Supplementary Figure 3.** Desmin (DES) and desmoplakin (DSP) immunostains of control (p.R406; top panels, Control line; bottom panels, IC-R406 line) and mutant (p.R406W; top panels, IM-R406W line; bottom panels, PT-R406W line) iPSC-CMs cultured on micropatterns. Scale bars, 20 µm.

#### Supplementary Table 1. Action potential parameters of hiPSC-CMs

**Spontaneous**

| | Control | | | IM-R406W | | | Mann-Whitney | |
|---|---|---|---|---|---|---|---|---|
| | Mean | ±sem | n | Mean | ±sem | n | Sign | p-value |
| Interval (ms) | 456.0 | 39.0 | 19 | 436.4 | 33.0 | 25 | ns | 0.7784 |
| MDP (mV) | -56.8 | 1.6 | 19 | -52.4 | 1.3 | 25 | * | 0.0335 |
| Overshoot (mV) | 34.2 | 2.1 | 19 | 41.4 | 1.9 | 25 | * | 0.0127 |
| Amplitude (mV) | 91.2 | 3.1 | 19 | 93.9 | 2.5 | 25 | ns | 0.4811 |
| dV/dt$_{max}$ (V/s) | 23.2 | 5.3 | 19 | 18.5 | 2.3 | 25 | ns | 0.3811 |
| APD$_{20}$ (ms) | 70.0 | 5.2 | 19 | 83.5 | 7.9 | 25 | ns | 0.3232 |
| APD$_{30}$ (ms) | 88.1 | 6.9 | 19 | 111.5 | 11.3 | 25 | ns | 0.2412 |
| APD$_{50}$ (ms) | 109.4 | 8.7 | 19 | 145.0 | 14.4 | 25 | ns | 0.1244 |
| APD$_{70}$ (ms) | 123.1 | 9.7 | 19 | 162.3 | 15.5 | 25 | ns | 0.1077 |
| APD$_{80}$ (ms) | 130.2 | 10.1 | 19 | 170.4 | 15.9 | 25 | ns | 0.1186 |
| APD$_{90}$ (ms) | 139.3 | 10.8 | 19 | 180.5 | 16.3 | 25 | ns | 0.1077 |

| | PT-R406W | | | IC-R406 | | | Mann-Whitney | |
|---|---|---|---|---|---|---|---|---|
| | Mean | ±sem | n | Mean | ±sem | n | Sign | p-value |
| Interval (ms) | 450.3 | 23.9 | 24 | 488.6 | 39.7 | 20 | ns | 0.7885 |
| MDP (mV) | -49.3 | 1.2 | 24 | -54.2 | 1.5 | 20 | ** | 0.0089 |
| Overshoot (mV) | 27.7 | 2.4 | 24 | 32.4 | 1.9 | 20 | ns | 0.2461 |
| Amplitude (mV) | 77.0 | 3.3 | 24 | 86.6 | 2.8 | 20 | ns | 0.0750 |
| dV/dt$_{max}$ (V/s) | 9.6 | 1.5 | 24 | 10.5 | 1.9 | 20 | ns | 0.3403 |
| APD$_{20}$ (ms) | 62.2 | 4.4 | 24 | 80.4 | 5.9 | 20 | * | 0.0383 |
| APD$_{30}$ (ms) | 76.4 | 5.6 | 24 | 103.8 | 8.5 | 20 | * | 0.0339 |
| APD$_{50}$ (ms) | 98.4 | 6.9 | 24 | 131.0 | 11.2 | 20 | * | 0.0431 |
| APD$_{70}$ (ms) | 114.6 | 8.5 | 24 | 149.3 | 12.5 | 20 | ns | 0.0674 |
| APD$_{80}$ (ms) | 124.4 | 9.1 | 24 | 158.6 | 13.1 | 20 | ns | 0.0921 |
| APD$_{90}$ (ms) | 137.7 | 9.9 | 24 | 170.8 | 13.9 | 20 | ns | 0.1018 |

**PCL1000**

| | Control | | | IM-R406W | | | Mann-Whitney | |
|---|---|---|---|---|---|---|---|---|
| | Mean | ±sem | n | Mean | ±sem | n | Sign | p-value |
| Interval (ms) | 1000.0 | 0.0 | 22 | 1000.0 | 0.0 | 15 | ns | 0.8115 |
| MDP (mV) | -91.5 | 0.1 | 22 | -91.8 | 0.4 | 15 | ns | 0.9452 |
| Overshoot (mV) | 46.0 | 3.1 | 22 | 52.3 | 3.1 | 15 | ns | 0.2132 |
| Amplitude (mV) | 137.5 | 3.1 | 22 | 144.1 | 3.0 | 15 | ns | 0.1612 |
| dV/dt$_{max}$ (V/s) | 88.1 | 15.5 | 22 | 77.2 | 10.8 | 15 | ns | 0.9149 |
| APD$_{20}$ (ms) | 65.3 | 7.9 | 22 | 96.5 | 11.3 | 15 | * | 0.0104 |
| APD$_{30}$ (ms) | 92.2 | 11.0 | 22 | 149.5 | 18.4 | 15 | ** | 0.0063 |
| APD$_{50}$ (ms) | 122.3 | 15.9 | 22 | 198.5 | 25.6 | 15 | ** | 0.0063 |
| APD$_{70}$ (ms) | 134.2 | 17.8 | 22 | 212.7 | 26.7 | 15 | ** | 0.0078 |
| APD$_{80}$ (ms) | 136.7 | 18.0 | 22 | 215.4 | 26.8 | 15 | ** | 0.0086 |
| APD$_{90}$ (ms) | 138.9 | 18.1 | 22 | 217.6 | 26.8 | 15 | ** | 0.0086 |

| | PT-R406W | | | IC-R406 | | | Mann-Whitney | |
|---|---|---|---|---|---|---|---|---|
| | Mean | ±sem | n | Mean | ±sem | n | Sign | p-value |
| Interval (ms) | 1000.0 | 0.0 | 30 | 1000.0 | 0.1 | 28 | ns | 0.0994 |
| MDP (mV) | -91.3 | 0.2 | 30 | -91.2 | 0.2 | 28 | ns | 0.9109 |
| Overshoot (mV) | 40.8 | 1.5 | 30 | 40.4 | 1.6 | 28 | ns | 0.7486 |
| Amplitude (mV) | 132.1 | 1.5 | 30 | 131.6 | 1.7 | 28 | ns | 0.787 |
| dV/dt$_{max}$ (V/s) | 65.6 | 13.9 | 30 | 57.3 | 7.8 | 28 | ns | 0.96 |
| APD$_{20}$ (ms) | 77.1 | 5.9 | 30 | 88.7 | 7.3 | 28 | ns | 0.2749 |
| APD$_{30}$ (ms) | 102.7 | 7.3 | 30 | 115.6 | 9.3 | 28 | ns | 0.3492 |
| APD$_{50}$ (ms) | 140.4 | 10.4 | 30 | 144.3 | 10.8 | 28 | ns | 0.7167 |
| APD$_{70}$ (ms) | 161.5 | 11.4 | 30 | 159.7 | 11.2 | 28 | ns | 0.9692 |
| APD$_{80}$ (ms) | 170.0 | 11.8 | 30 | 165.8 | 11.5 | 28 | ns | 0.9537 |
| APD$_{90}$ (ms) | 179.7 | 12.2 | 30 | 174.0 | 12.4 | 28 | ns | 0.9323 |

**PCL700**

| | Control | | | IM-R406W | | | Mann-Whitney | |
|---|---|---|---|---|---|---|---|---|
| | Mean | ±sem | n | Mean | ±sem | n | Sign | p-value |
| Interval (ms) | 700.0 | 0.0 | 21 | 700.0 | 0.0 | 15 | ns | 0.582 |
| MDP (mV) | -91.4 | 0.2 | 21 | -91.4 | 0.3 | 15 | ns | 0.9243 |
| Overshoot (mV) | 46.2 | 2.9 | 21 | 51.7 | 3.0 | 15 | ns | 0.2651 |
| Amplitude (mV) | 137.5 | 2.8 | 21 | 143.1 | 2.8 | 15 | ns | 0.2937 |
| dV/dt$_{max}$ (V/s) | 96.3 | 17.9 | 21 | 80.0 | 11.2 | 15 | ns | >0.9999 |
| APD$_{20}$ (ms) | 61.2 | 8.8 | 21 | 87.4 | 11.1 | 15 | * | 0.026 |
| APD$_{30}$ (ms) | 85.5 | 10.9 | 21 | 128.5 | 16.2 | 15 | * | 0.0389 |
| APD$_{50}$ (ms) | 112.8 | 14.5 | 21 | 168.6 | 21.3 | 15 | * | 0.0303 |
| APD$_{70}$ (ms) | 124.3 | 15.9 | 21 | 184.0 | 21.7 | 15 | * | 0.0255 |
| APD$_{80}$ (ms) | 126.9 | 16.1 | 21 | 187.2 | 21.7 | 15 | * | 0.0233 |
| APD$_{90}$ (ms) | 129.2 | 16.2 | 21 | 189.8 | 21.6 | 15 | * | 0.0233 |

| | PT-R406W | | | IC-R406 | | | Mann-Whitney | |
|---|---|---|---|---|---|---|---|---|
| | Mean | ±sem | n | Mean | ±sem | n | Sign | p-value |
| Interval (ms) | 699.9 | 0.1 | 27 | 700.0 | 0.1 | 25 | ns | 0.28 |
| MDP (mV) | -91.1 | 0.1 | 27 | -91.3 | 0.2 | 25 | ns | 0.2236 |
| Overshoot (mV) | 40.8 | 1.6 | 27 | 39.7 | 1.7 | 25 | ns | 0.6238 |
| Amplitude (mV) | 131.9 | 1.6 | 27 | 131.0 | 1.8 | 25 | ns | 0.7029 |
| dV/dt$_{max}$ (V/s) | 62.4 | 10.6 | 27 | 61.1 | 10.1 | 25 | ns | 0.8881 |
| APD$_{20}$ (ms) | 73.0 | 6.1 | 27 | 79.2 | 7.1 | 25 | ns | 0.467 |
| APD$_{30}$ (ms) | 96.3 | 7.4 | 27 | 102.3 | 8.5 | 25 | ns | 0.611 |
| APD$_{50}$ (ms) | 130.8 | 10.0 | 27 | 129.0 | 9.7 | 25 | ns | 0.9566 |
| APD$_{70}$ (ms) | 152.0 | 11.1 | 27 | 144.2 | 10.3 | 25 | ns | 0.7889 |
| APD$_{80}$ (ms) | 160.4 | 11.4 | 27 | 150.1 | 10.8 | 25 | ns | 0.5983 |
| APD$_{90}$ (ms) | 170.7 | 11.9 | 27 | 158.2 | 12.0 | 25 | ns | 0.5012 |

Test: Unpaired non-parametric t-test (Mann-Whitney)
Control vs IM-R406W
PT-R406W vs IC-R406

ns = not significant
*: $p < 0.05$
**: $p < 0.01$

**Supplementary Table 2. Action potential parameters of impaled EHTs**

Spontaneous

| | Control | | | IM-R406W | | | |
|---|---|---|---|---|---|---|---|
| | Mean | ± sem | n | Mean | ± sem | n | Sign |
| Interval (s) | 1.7 | 0.1 | 6 | 1.4 | 0.2 | 4 | ns |
| MDP (mV) | -66.2 | 1.6 | 6 | -71.4 | 5.9 | 4 | ns |
| Overshoot (mV) | 33.5 | 2.4 | 6 | 35.8 | 3.2 | 4 | ns |
| Amplitude (mV) | 100.3 | 4.1 | 6 | 106.1 | 2.5 | 4 | ns |
| $dV/dt_{max}$ (V/s) | 77.6 | 18.1 | 6 | 116.5 | 27.6 | 4 | ns |
| $APD_{20}$ (ms) | 118.8 | 5.7 | 6 | 192.7 | 25.0 | 4 | * |
| $APD_{30}$ (ms) | 149.0 | 6.6 | 6 | 244.0 | 33.1 | 4 | ** |
| $APD_{50}$ (ms) | 177.8 | 8.7 | 6 | 284.3 | 37.1 | 4 | * |
| $APD_{70}$ (ms) | 194.6 | 10.5 | 6 | 299.9 | 38.0 | 4 | * |
| $APD_{80}$ (ms) | 205.0 | 11.1 | 6 | 308.0 | 38.3 | 4 | * |
| $APD_{90}$ (ms) | 229.8 | 12.3 | 6 | 323.4 | 37.6 | 4 | ns |

| | PT-R406W | | | IC-R406 | | | |
|---|---|---|---|---|---|---|---|
| | Mean | ± sem | n | Mean | ± sem | n | Sign |
| Interval (s) | 0.8 | 0.0 | 7 | 1.0 | 0.2 | 4 | ns |
| MDP (mV) | -73.3 | 4.6 | 7 | -67.1 | 2.7 | 4 | ns |
| Overshoot (mV) | 30.5 | 2.8 | 7 | 33.1 | 0.4 | 4 | ns |
| Amplitude (mV) | 100.6 | 5.0 | 7 | 102.1 | 2.3 | 4 | ns |
| $dV/dt_{max}$ (V/s) | 36.8 | 9.5 | 7 | 14.6 | 1.8 | 4 | ns |
| $APD_{20}$ (ms) | 196.7 | 15.7 | 7 | 150.8 | 9.6 | 4 | ns |
| $APD_{30}$ (ms) | 254.6 | 20.3 | 7 | 193.8 | 16.8 | 4 | ns |
| $APD_{50}$ (ms) | 310.1 | 22.8 | 7 | 243.8 | 32.7 | 4 | ns |
| $APD_{70}$ (ms) | 332.5 | 23.8 | 7 | 269.8 | 40.6 | 4 | ns |
| $APD_{80}$ (ms) | 342.3 | 24.2 | 7 | 282.9 | 43.6 | 4 | ns |
| $APD_{90}$ (ms) | 355.0 | 25.0 | 7 | 302.6 | 48.9 | 4 | ns |

Test: Unpaired non-parametric t-test (Mann-[illegible]
Control vs IM-R406W
PT-R406W vs IC-R406

ns = not significant
*: $p < 0.05$
**: $p < 0.01$

**Supplementary Table 3. Omics data**

https://uncloud.univ-nantes.fr/index.php/s/tRyQmy9tgkXHzDw

**Supplementary table 4. ECG parameters in 10- and 20-week-old wildtype (WT) and Des-p.R405W knock-in (KI) female and male mice under baseline conditions.**

| | **RR interval (ms)** | **P wave duration (ms)** | **PR interval (ms)** | **QRS duration (ms)** | **QT interval (ms)** | **R wave ampl. (mV)** | **S wave ampl. (mV)** |
|---|---|---|---|---|---|---|---|
| ***10 weeks old*** | | | | | | | |
| Female WT (24) | 121.8 ± 12.1 | 13.0 ± 1.9 | 38.8 ± 3.5 | 10.8 ± 0.9 | 53.0 ± 4.39 | 1.1 ± 0.2 | -0.3 ± 0.2 |
| Female KI (18) | 127.5 ± 13.8 | 13.4 ± 2.0 | 38.5 ± 2.6 | 11.3 ± 0.7 [$] | 48.2 ± 3.4 | 0.9 ± 0.3 [$] | -0.3 ± 0.1 |
| Male WT (16) | 125.8 ± 9.4 | 13.0 ± 1.5 | 37.4 ± 2.3 | 11.1 ± 0.9 | 53.5 ± 5.0 | 0.9 ± 0.2 | -0.3 ± 0.2 |
| Male KI (21) | 126.0 ± 9.9 | 14.0 ± 1.3 * | 38.5 ± 2.9 | 11.9 ± 1.5 [$] | 48.3 ± 4.1 | 0.8 ± 0.2 * | -0.3 ± 0.2 |
| ***20 weeks old*** | | | | | | | |
| Female WT (15) | 122.2 ± 9.8. | 13.0 ± 2.2 | 39.0 ± 2.5 | 10.7 ± 1.1 | 52.9 ± 3.7 | 0.9 ± 0.2 | -0.2 ± 0.1 |
| Female KI (9) | 126.8 ± 12.1 | 13.0 ± 2.3 | 40.8 ± 1.5 | 11.1 ± 1.1 | 53.1 ± 3.7 | 0.8 ± 0.2 | -0.4 ± 0.1 * |
| Male WT (10) | 130.5 ± 13.1 | 13.1 ± 2.2 | 38.1 ± 3.5 | 10.3 ± 0.5 | 50.9 ± 2.0 | 0.8 ± 0.2 | -0.2 ± 0.1 |
| Male KI (10) | 132.0 ± 16.9 | 12.5 ± 1.3 | 38.3 ± 2.0 | 12.0 ± 1.6 ** | 53.2 ± 4.2 | 0.7 ± 0.2 | -0.2 ± 0.1 |

Abbreviation: ampl., amplitude. Number of mice within parentheses. Data are expressed as mean ± standard deviation. *, **: $p < 0.05$ and $p < 0.01$, respectively, *versus* WT at corresponding age (Student t-test or Mann-Whitney test). [§] $0.05 > p < 0.07$, *versus* WT.

**Supplementary table 5. Heart weight to body weight (HW/BW) and heart weight to tibia length (HW/TL) ratios in female and male wildtype (WT) and Des-p.R405W knock-in (KI) mice at the age of 11 and 21 weeks.**

| **Age** | **11 weeks** | | | | **21 weeks** | | | |
|---|---|---|---|---|---|---|---|---|
| **Sex** | **Female** | | **Male** | | **Female** | | **Male** | |
| **Genotype (n)** | **WT (8)** | **KI (7)** | **WT (3)** | **KI (6)** | **WT (15)** | **KI (7)** | **WT (8)** | **KI 12)** |
| **HW/BW** | 5.39 ± 0.49 | 5.44 ± 0.48 | 5.22 ± 0.18 | 5.31 ± 0.41 | 5.33 ± 0.61 | 5.52 ± 0.56 | 6.19 ± 0.96 | 5.81 ± 0.91 |
| **HW/TL** | 6.76 ± 0.49 | 6.67 ± 0.49 | 8.11 ± 0.67 | 8.49 ± 0.84 | 7.33 ± 0.67 | 8.05 ± 0.69 * | 10.39 ± 1.27 | 10.52 ± 1.23 |

Values are means ± standard deviation. *, $p < 0.05$ *versus* WT at corresponding age (Mann-Whitney test).

**Supplementary table 6. Percentage of fibrosis in ventricular sections of 10 wildtype and 10 Des-p.R405W knock-in mice at the age of 21 weeks.**

| | Right ventricle | Septum | Left ventricle |
|---|---|---|---|
| **Wildtype mice** | 2.38 ± 1.13 | 1.10 ± 0.35 *** | 1.52 ± 0.66 * |
| **Knock-in mice** | 2.82 ± 1.08 | 1.02 ± 0.65 *** | 1.49 ± 0.25 *** |

Values are means ± standard deviation. *, ***: $p < 0.05$ and $p < 0.001$, respectively, *versus* corresponding value in the right ventricle.
